# All-optical reporting of inhibitory receptor driving force in the nervous system

**DOI:** 10.1101/2023.08.30.555464

**Authors:** Joshua S. Selfe, Teresa J. S. Steyn, Eran F. Shorer, Richard J. Burman, Kira M. Düsterwald, Ahmed S. Abdelfattah, Eric R. Schreiter, Sarah E. Newey, Colin J. Akerman, Joseph V. Raimondo

## Abstract

Ionic driving forces provide the net electromotive force for ion movement across receptors, channels, and transporters, and are a fundamental property of all cells. In the brain for example, fast synaptic inhibition is mediated by chloride permeable GABAA receptors, and single-cell intracellular recordings have been the only method for estimating driving forces across these receptors (DFGABAA). Here we present a new tool for quantifying inhibitory receptor driving force named ORCHID: all-Optical Reporting of CHloride Ion Driving force. We demonstrate ORCHID’s ability to provide accurate, high-throughput measurements of resting and dynamic DFGABAA from genetically targeted cell types over multiple timescales. ORCHID confirms theoretical predictions about the biophysical mechanisms that establish DFGABAA, reveals novel differences in DFGABAA between neurons and astrocytes, and affords the first *in vivo* measurements of intact DFGABAA. This work extends our understanding of inhibitory synaptic transmission and establishes a precedent for all-optical methods to assess ionic driving forces.

## Introduction

Ionic driving forces represent the net electromotive force available for generating ion fluxes across receptors, channels and transporters. A driving force is the product of both an ion’s equilibrium potential and the potential difference across the membrane, and constitutes the energy per unit charge that is available to mediate a host of fundamental cellular processes^1^. Despite their fundamental importance, the only way to estimate ionic driving forces has been to perform intracellular electrophysiological recording methods. This has limited measurements to small numbers of cells and, more fundamentally, in all cases these methods either directly, or indirectly, affect the ion gradients that underlie the driving forces^2^. Meanwhile, methods for imaging intracellular ions cannot provide information on ionic driving force because they are blind to the cell’s membrane potential. Here we describe the first all-optical strategy for measuring ionic driving forces and demonstrate the utility of this approach for performing rapid, multi-cell measurements of undisturbed driving forces.

In the nervous system, fast synaptic inhibition is mediated by anion-permeable type A γ-aminobutyric acid receptors (GABAARs) and glycine receptors (GlyRs). These ligand-gated receptors are primarily permeable to chloride (Cl^-^) and, to a lesser extent, bicarbonate (HCO3-) ^3^. The transmembrane gradients for Cl^-^ and HCO3-determine the inhibitory receptor reversal potential (EGABAA) and, together with the membrane potential (Vm), set the ionic driving force across these receptors (e.g. DFGABAA). DFGABAA, by controlling anion flux across inhibitory receptors, is therefore a key factor in determining inhibitory synaptic signaling in the nervous system. As an exemplar of an all-optical strategy for measuring ionic driving forces, we present ORCHID (all-Optical Reporting of CHloride Ion Driving force) as a new tool for quantifying DFGABAA, which is based on combining genetically encoded voltage indicators with light-gated ion channels

Intracellular Cl^-^ concentration ([Cl^-^]i) and hence EGABAA, is a dynamic variable that is known to differ between subcellular compartments, cell types, and over a range of timescales and network states^4^. Owing to the fact that EGABAA is typically close to Vm, relatively small changes to EGABAA or Vm can have a large effect upon the polarity and magnitude of DFGABAA, and thus upon inhibitory signalling^5^. Alterations to neuronal DFGABAA have been implicated in multiple neurological diseases including epilepsy, chronic pain, schizophrenia, and autism^6–9^. Similarly, DFGABAA is thought to be modulated during brain development as part of the GABAergic system’s contribution to neural circuit formation^10,11^. In regions of the mature nervous system, it is believed that DFGABAA varies diurnally and is regulated as a function of wakefulness in neurons^12,13^ and astrocytes^14^. Therefore, given the central importance of DFGABAA for both brain function and dysfunction, techniques for estimating DFGABAA are of tremendous interest to the neuroscience field.

The most widely used electrophysiological strategy for estimating DFGABAA is gramicidin-perforated patch clamp recording^2^. This technique provides electrical access to a target cell of interest without directly disturbing [Cl^-^]i. When combined with a means of generating ion selective conductances, such as activation of GABAARs or light-sensitive channels with similar Cl^-^ permeability^15^, EGABAA, and DFGABAA can be estimated. However, gramicidin recordings are technically challenging, have a low success rate and throughput, and perturb the intracellular levels of HCO3-, directly modifying DFGABAA, as well as K^+^ and Na^+^, which can indirectly change [Cl^-^]i and DFGABAA via the action of cation-Cl^-^ cotransporters (CCCs) such as KCC2 and NKCC1^1^. Meanwhile, the best optical strategies for [Cl^-^]i estimation include fluorescence lifetime imaging (FLIM) of the Cl^-^-sensitive dye MQAE^16,17^ and genetically encoded Cl^-^ indicators^18^. Ratiometric genetically encoded Cl^-^ indicators allow for high-throughput measurements and can be targeted to different cell types, although there are challenges associated with ion selectivity, signal-to-noise ratio, and measurements in intact systems^19–22^. More fundamentally however, whilst currently available optical strategies can provide an estimate of [Cl^-^]i, they do not incorporate information about the cell’s Vm, and therefore cannot quantify the force available in the transmembrane gradient for Cl^-^ and HCO3-, which is DFGABAA.

As we show below, ORCHID provides easy-to-use, genetically encoded optical estimation of a cell’s undisturbed DFGABAA. ORCHID has been designed to incorporate the sensitive, chemi-genetically encoded voltage indicator (GEVI), Voltron2-ST^23^, with an independent means of generating anion selective currents by using light activation of *Guillardia theta* anion channelrhodopsin 2 (GtACR2)^24^. GtACR2’s anion permeability follows the lyotropic series characteristic of GABAARs and GlyRs^24–26^ and the reversal potentials for GtACR2 and GABAARs are equivalent^27,28^. This means that they share the same relative permeability to Cl^-^ and HCO3-. Voltron2-ST is therefore used to measure the magnitude and direction of GtACR2 anion-current-induced changes to Vm, thereby providing an estimate of DFGABAA. We initially validate the ORCHID approach in mice by demonstrating that it replicates estimates of DFGABAA that rely upon the activation of endogenous GABAAR conductances. ORCHID is shown to provide reliable measurements of resting and dynamic DFGABAA in different genetically targeted cell populations and across a timescale of seconds and hours. ORCHID is also shown to confirm theoretical predictions regarding the biophysical mechanisms that establish DFGABAA. Finally, we use ORCHID to reveal novel cell-type-specific differences in activity-dependent DFGABAA changes during periods of network activity, and to provide the first *in vivo* measurements of intact DFGABAA in mouse cortical neurons. Together, our results demonstrate that ORCHID allows high-throughput, dynamic estimation of DFGABAA in a cell-type-specific manner. This will assist in the study of inhibitory synaptic transmission and Cl^-^ homeostasis in the nervous system, and establishes a precedent for using all-optical methods to assess ionic driving force more generally.

## Results

### ORCHID provides an optical estimate of the magnitude and direction of inhibitory receptor driving force

In order to optically measure DFGABAA, we designed a strategy based on combining a genetically encoded optical reporter of Vm, with the simultaneous activation of a light-sensitive transmembrane anion channel (**Fig. 1a-b**). The resulting ORCHID strategy utilizes the recently developed soma-targeted GEVI Voltron2 (Voltron2-ST), and the light-activated anion channel GtACR2, separated via a P2A linker sequence (**Fig. 1c**). The ORCHID construct has a double-floxed design for targeting Cre-expressing cells and is under the control of the EF1α promoter (see **Supplementary Fig. 1** for construct design). Cre-expressing neurons in mouse hippocampal organotypic brain slices were successfully transduced with adeno-associated virus (AAV) vectors encoding ORCHID (see **Methods**). Prior to imaging, a dye incubation step results in Voltron2-ST protein being irreversibly bound by one of several possible Janelia Fluor (JF) dyes, each of which has different spectral properties^29^.

**Figure 1:**
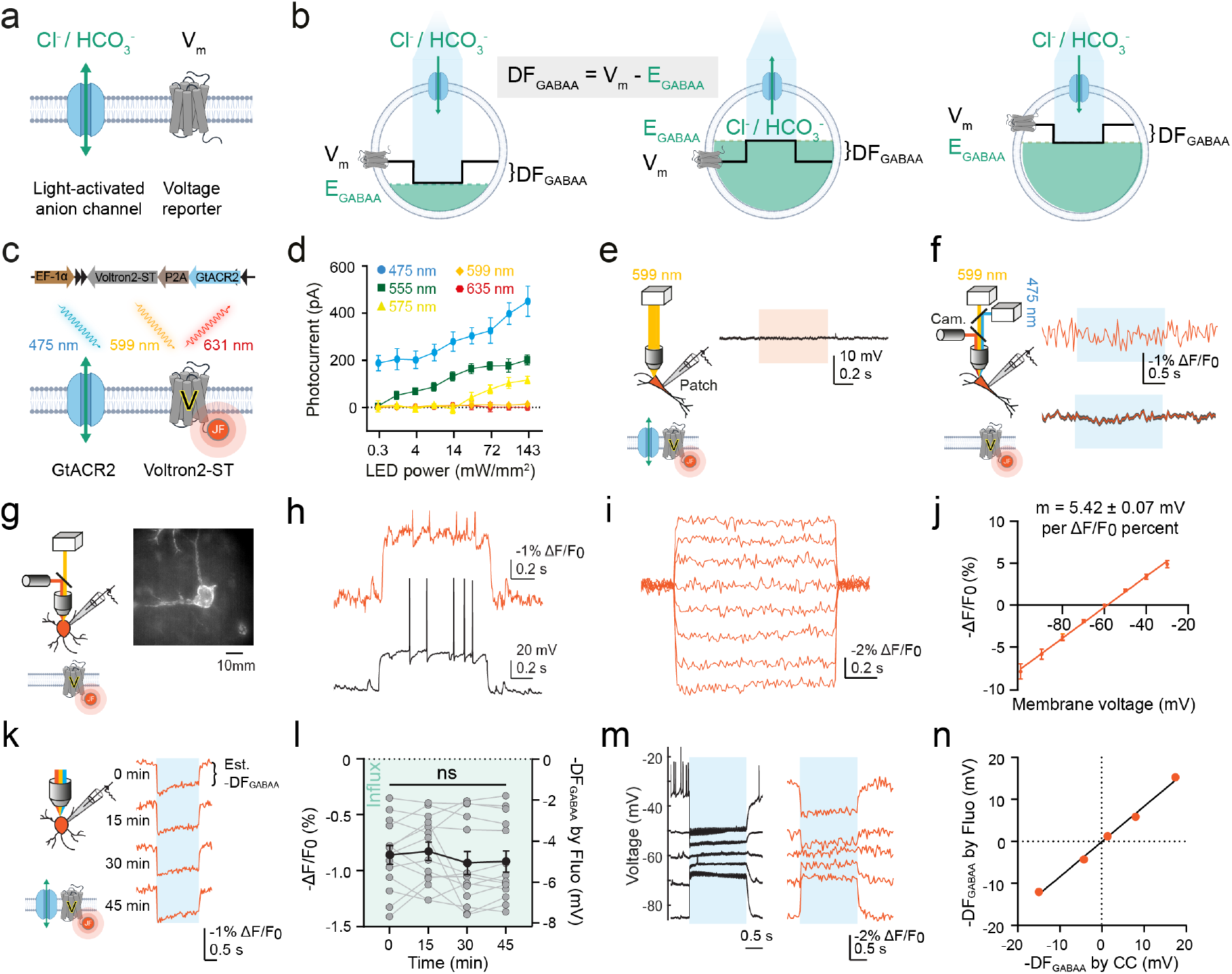
ORCHID provides an optical estimate of the magnitude and direction of inhibitory receptor driving force. **a**, Conceptual design of an all-optical strategy to estimate DFGABAA, combining a light-activated anion channel with an optical voltage reporter. **b**, Activation of an anion conductance (blue) drives Vm (black trace) toward EGABAA (green dotted line). The Vm shift from before to after anion channel activation provides an estimate of DFGABAA (DFGABAA = Vm – EGABAA). Left, EGABAA is negative with respect to initial Vm, constituting a driving force that results in anion influx and hyperpolarization. Middle, due to higher [Cl^-^]i (green shading), EGABAA is positive with respect to initial Vm, and DFGABAA now causes anion efflux and depolarization. Right, despite identical EGABA as in ‘middle’, initial Vm results in a DFGABAA that causes influx and hyperpolarization **c**, ORCHID utilizes co-expression of GtACR2 with Voltron2-ST and JF608 dye. **d**, Whole-cell patch clamp recorded GtACR2 photocurrents at different wavelengths from hippocampal neurons. **e**, Current-clamp recording from an ORCHID expressing neuron demonstrates no change in Vm during orange light exposure (orange rectangle). **f**, Voltage imaging of a hippocampal neuron expressing Voltron2608-ST alone (no GtACR2) reveals that blue light delivery strobed between camera exposures (see Methods) causes no change in measured fluorescence (top, orange trace); averaged recorded fluorescence from 29 neurons showing mean ± standard error of the mean (SEM; bottom, orange and grey respectively). **g**, Setup for patch-clamp characterization of Voltron2608-ST. Inset: widefield image of neuron patched in ‘h’; scale bar: 10 µm. **h**, Current-clamp recording (black trace) and Voltron2608-ST fluorescence response (orange trace) following a 200 pA current injection. **i**, Voltron2608-ST fluorescence response to 10 mV voltage steps (Vhold = -60 mV). **j**, Voltage-fluorescence relationship of Voltron2608-ST, with a slope 5.42 ± 0.07 mV per ΔF/F0 percent (R^2^ = 0.9311, *n* = 7). **k**, DFGABAA estimated using fluorescence transients (orange traces) of an ORCHID expressing neuron following blue light activation (blue rectangle) was stable over 45 min. **l**, Population data showing stable estimates of DFGABAA using ORCHID (repeated measures one-way ANOVA, *P* = 0.2469, *n* = 12). Green shading indicates direction of anion flux. **m**, ORCHID used to estimate DFGABAA in neurons current clamped at a range of Vm values (left panel, Vm; right panel, fluorescence). Action potentials are truncated due to averaging. **n**, Fluorescence-voltage relationship of Voltron2608-ST (j) was used to convert ΔF/F0 measurements at each initial Vm to DFGABAA values, which were equivalent to DFGABAA recorded using current-clamp recordings (R^2^ = 0.9955). ns = not significant (*P* > 0.05); error bars indicate mean ± SEM.

For ORCHID to work, it is critical to achieve spectral separation of the voltage readout using Voltron2-ST, from the manipulation of anion currents using GtACR2. Whole-cell patch-clamp recordings from hippocampal pyramidal neurons expressing GtACR2 revealed a wide activation spectrum, with currents being generated by blue (475/28 nm), green (555/28 nm), and yellow (575/25 nm) excitation wavelengths, but not by orange (599/13 nm) or red (635/22 nm) wavelengths (**Fig. 1d**). We therefore utilized JF608 together with orange (599/13 nm) excitation light to ensure that GEVI imaging elicited no GtACR2 currents (**Fig. 1e**). Activation of GtACR2 with blue light (475/28 nm) does not significantly excite JF608 fluorescence^30^; nonetheless, blue light delivery was rapidly interleaved with camera acquisition to ensure that blue-light-generated autofluorescence would not cause artefacts in the orange GEVI imaging channel (see **Methods**). Imaging neurons expressing only Voltron2-ST labelled with JF608 (Voltron2608-ST) confirmed that there was no change in fluorescence during blue light delivery (**Fig. 1f**). Meanwhile, whole-cell patch-clamp recordings from ORCHID-expressing neurons confirmed that Voltron2608-ST reports linear fluorescence changes in response to voltage steps, thereby establishing that ΔF/F0 can be used to estimate changes in voltage (ΔV) in subsequent analyses (**Fig. 1g-j**). In ORCHID-expressing neurons, blue light stimulation elicited clear and reproducible Vm shifts, the polarity and magnitude of which could be used to estimate DFGABAA, and which showed excellent reproducibility within the same neuron (**Fig. 1k-l**).

To assess ORCHID’s ability to quantify DFGABAA, we performed simultaneous current-clamp recordings and voltage imaging in ORCHID-expressing neurons. Different DFGABAA values were imposed by injecting current via the patch pipette. Meanwhile, voltage imaging of ORCHID was performed under orange light, and anion currents were generated via blue-light activation of ORCHID (**Fig. 1m-n**). The relationship between ΔF/F0 and ΔV (**Fig. 1j**) was used to convert the measured ΔF/F0 values into estimations of anion-current-induced changes to Vm; these optically determined DFGABAA values were found to be equivalent to DFGABAA measured from the simultaneous current-clamp recordings (**Fig. 1n**; R^2^ = 0.9955). This demonstrated that the ability of ORCHID to estimate DFGABAA is as accurate as using current-clamp recordings, with both methods accurately determining the direction of DFGABAA and inclined to underestimate the full magnitude of DFGABAA. ORCHID’s ability to detect a change in DFGABAA was further confirmed by comparing measurements of DFGABAA made before and after a neuron experienced an increase in [Cl^-^]i imposed by dialysis (**Supplementary Fig. 2**).

As further validation, we assessed ORCHID’s ability to accurately determine DFGABAA when the anion currents were generated by activating endogenous GABAARs in different neuronal populations. Transgenic mice expressing Cre recombinase in different cell types were used to restrict reporter expression to either CaMKIIα+ pyramidal neurons or GAD2+ interneurons. Voltron2-ST labelled with JF549 (Voltron2549-ST) was used to assess DFGABAA via endogenous GABAARs on the cell soma, which were activated by picolitre delivery of GABA (500 µM) via a glass micropipette and in the presence of a GABABR antagonist (5 µM CGP-55845). This was found to be an effective method of estimating DFGABAA (**Supplementary Fig. 3, 4 and 5**) and revealed that both CaMKIIα+ pyramidal neurons and GAD2+ interneurons exhibit hyperpolarizing DFGABAA under these conditions, with GAD2+ interneurons exhibiting slightly greater DFGABAA (**Fig. 2a-c**). When these experiments were conducted using ORCHID, our all-optical strategy revealed similar hyperpolarizing DFGABAA values for both neuronal populations, again with slightly greater DFGABAA in GAD2+ interneurons compared to CaMKIIα+ pyramidal neurons (**Fig. 2d-f**). DFGABAA estimates made using GtACR2 versus endogenous GABAARs were not statistically distinguishable (CaMKIIα+ pyramidal neurons: Mann-Whitney test, *P* = 0.066; GAD2+ interneurons: Mann-Whitney test, *P* = 0.1819).

**Figure 2:**
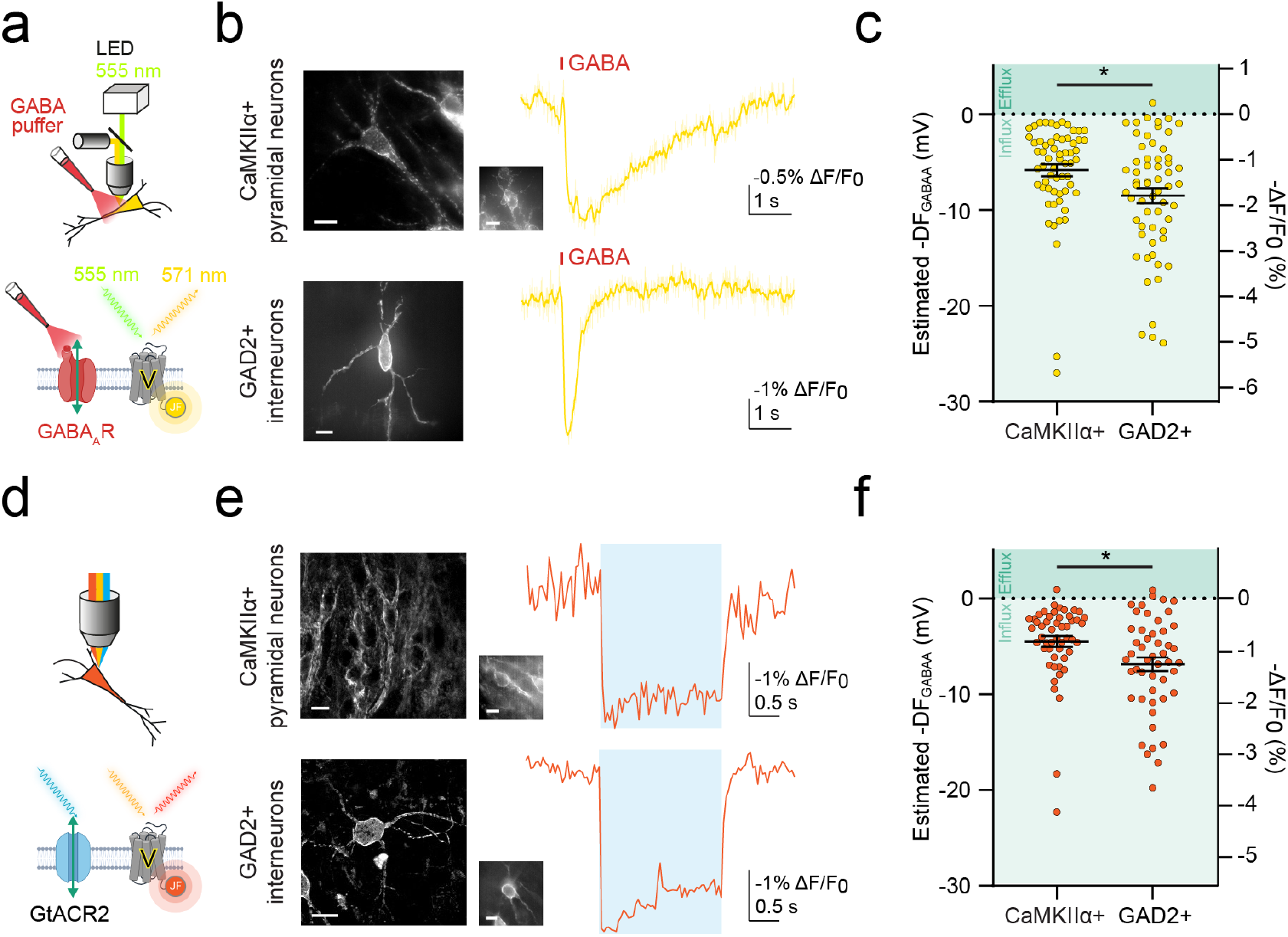
ORCHID reports endogenous inhibitory receptor driving force in different neuronal populations. **a**, Schematics showing DFGABAA estimation by imaging Voltron2549-ST together with activation of endogenously expressed GABAARs via soma-directed GABA application (red; 500 µM). CGP-55845 (5 µM) was used to block GABABRs. **b**, Widefield fluorescence images of CaMKIIα+ pyramidal neurons (top) and GAD2+ interneurons (bottom) expressing Voltron2549-ST (left) and fluorescence recordings used to estimate DFGABAA from each cell type (right, yellow traces), with recorded cells shown via inset if not in the main image; scale bar: 10 µm. Red bar indicates GABA application. **c**, Population data showing estimated DFGABAA was different between cell types (CaMKIIα+: 5.85 ± 0.62 mV vs GAD2+: 8.52 ± 0.79 mV, Mann-Whitney test, *P* = 0.0068, *n* = 62 and 60 cells respectively). Green shading indicates direction of anion flux. **d**, Schematics depicting ORCHID’s use for estimating DFGABAA in neurons. **e**, Left, confocal images of CaMKIIα+ pyramidal neurons (top) and GAD2+ interneurons (bottom) expressing ORCHID in mouse organotypic hippocampal brain slices. Right, fluorescence recordings used to estimate DFGABAA from each cell type, with widefield images of the recorded cells (inset); scale bar: 10 µm. **f**, Estimated DFGABAA was significantly different between cell types (CaMKIIα+: 4.52 ± 0.57 mV vs GAD2+: 6.88 ± 0.72 mV, Mann-Whitney test, *P* = 0.0057, *n* = 51 and 49 cells respectively). ns = not significant (*P* > 0.05); \**P* ≤ 0.05; error bars indicate mean ± SEM.

### ORCHID strategies confirm theoretical predictions regarding inhibitory receptor driving force

The accepted view is that DFGABAA is predominantly established by the action of CCCs such as KCC2 and NKCC1, which are able to use secondary active transport to move Cl^-^ against its transmembrane concentration gradient by utilizing cation gradients established by the primary active transporter, Na⁺/K⁺-ATPase^1^. However, work using the genetically encoded Cl^-^ indicator, Clomeleon, challenged this idea by proposing that impermeant anions (and not CCCs) set [Cl^-^]i, EGABAA and DFGABAA31. We sought to examine these biophysical principles by first using a theoretical model to make predictions about the mechanisms that establish DFGABAA. We extended a computational biophysical model of a single neuron based on the pump-leak mechanism that incorporated the permeant ions (Na^+^, K^+^, Cl^-^, HCO3-) and their movement via leak channels, receptors, the Na^+^/K^+^-ATPase, and the major Cl^-^ extruder in mature neurons, KCC2. The model also incorporated impermeant anions, X, with average charge z^32^, pH control via the HCO3--buffering system, plus Na^+^/H^+^ exchange, dynamic volume, water permeability, and synaptic GABAARs that were able to elicit Cl^-^ and HCO3-conductances and resulting changes in Vm (**Fig. 3a**). As expected, simulating a block of KCC2 in the model caused a positive shift in both ECl, EGABAA and [Cl^-^]i, with a small effect on Vm, (**Fig. 3b**). This resulted in a substantial shift in DFGABAA causing a decrease in amplitude and an inversion in polarity of GABAAR-mediated potentials (**Fig. 3c**). Meanwhile, simulating the addition of impermeant anions by reducing their average charge z generated a persistent negative shift in ECl, EGABAA and [Cl^-^]i, consistent with previous observations^32^. However, this manipulation caused ionic redistribution such that Vm also shifted and in a manner proportional to the effects upon ECl and EGABAA (**Fig. 3d**). Therefore GABAAR-mediated potentials were of a similar size following the addition of impermeant anions (**Fig. 3e**).

**Figure 3:**
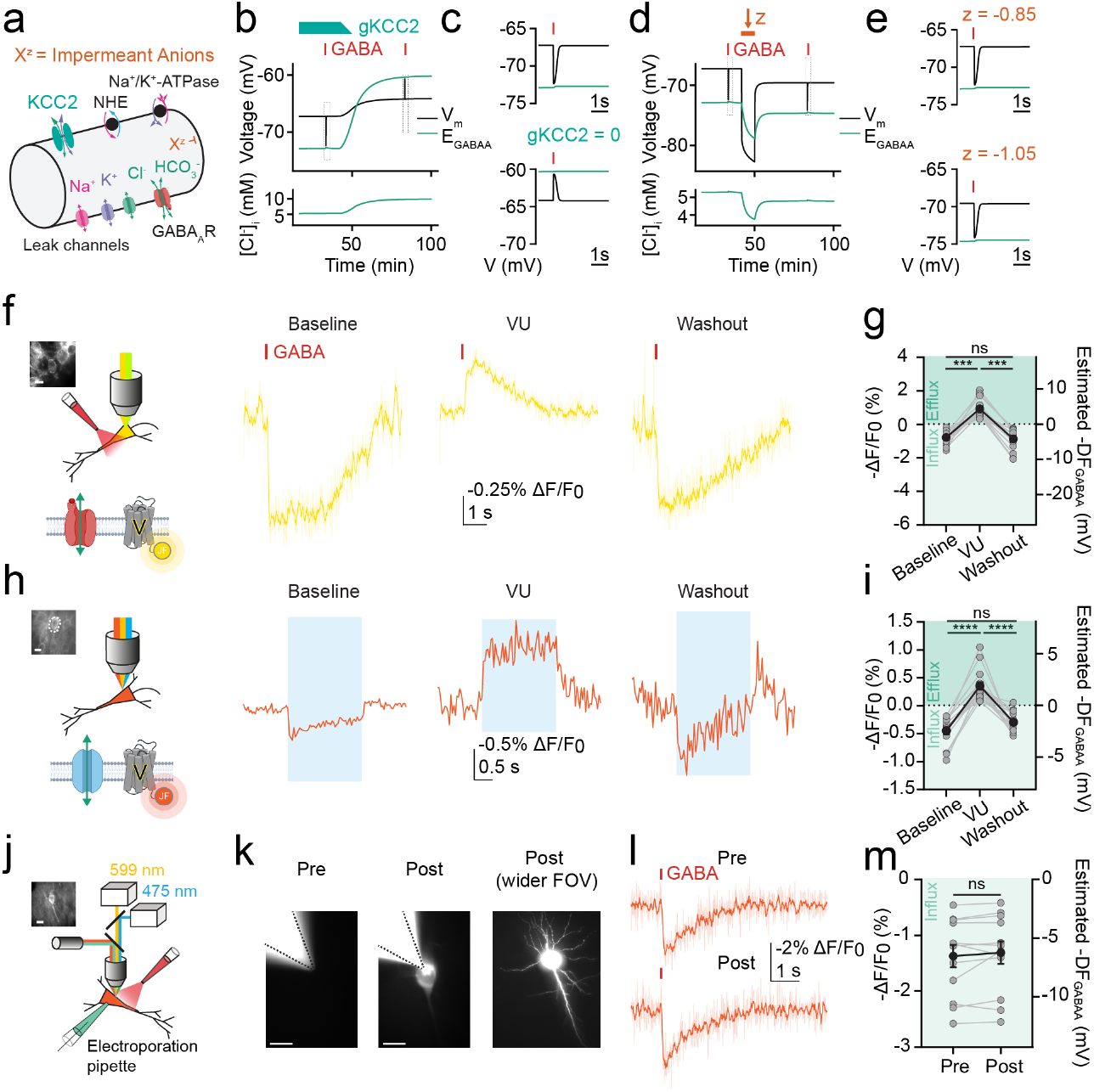
ORCHID strategies confirm theoretical predictions regarding inhibitory receptor driving force. **a**, Biophysical computational model of ion dynamics based on the pump-leak mechanism. Transmembrane movement of permeant ions (Na^+^, K^+^, Cl^-^, HCO3-) was modelled though leak channels, KCC2, Na^+^/K^+^-ATPase, and GABAARs. Impermeant anions, X, with average charge z could not cross the membrane. **b**, Top, simulated decrease in gKCC2 from 20 µS/cm^2^ to 0 µS/cm^2^ between 41 min and 50 min (teal bar) caused an increase in EGABAA (green) and smaller changes in Vm (black). Simulated synaptic GABAAR activation (red bar and grey rectangle) delivered before and after simulated KCC2 blockade. Bottom, increased [Cl^-^]i following gKCC2 reduction. **c**, Enlarged view of grey rectangles in ‘b’ show simulated synaptic GABAAR activation (red bar) in the neuron causes hyperpolarization with KCC2 intact (top), but depolarization with KCC2 blocked (bottom), indicating a shift from a hyperpolarizing to depolarizing driving force. **d,** As in ‘b’ but with average intracellular impermeant anion charge (z) shifted from -0.85 to -1.05 between 41 min and 50 min (orange bar). Top, proportional drop in Vm (black) and EGABAA (green) reflected a relatively unchanged DFGABAA despite altered z. Bottom, reduced [Cl^-^]i in response to decreased z. **e**, Enlarged view of grey rectangles in ‘d’ show simulated synaptic GABAAR activation results in similarly sized hyperpolarization in the neuron with default z of -0.85 (top), and with reduced z of -1.05 (bottom). **f**, Voltage imaging (Voltron2549-ST, yellow traces) of a CaMKIIα+ pyramidal neuron (inset, scale bar: 10 µm) following activation of endogenous GABAARs at baseline (left), during KCC2 blockade by VU (middle; 10 μM), and after VU washout (right). **g**, Population data showing blockade of KCC2 by VU caused a significant depolarization of DFGABAA in CaMKIIα+ neurons (left, Wilcoxon matched-pairs signed rank tests, wash-in *P* = 0.0001, washout *P* = 0.0001, *n* = 14, baseline vs washout comparison, *P* = 0.6698). Green shading indicates direction of anion flux. **h**, All-optical ORCHID was used to record DFGABAA as in ‘f’. **i**, Population data showing VU caused a significant shift in DFGABAA recorded using ORCHID in CaMKIIα+ pyramidal neurons (left, Wilcoxon matched-pairs signed rank tests, wash-in *P* < 0.0001, washout *P* < 0.0001, *n* = 16, baseline vs washout comparison, *P* = 0.1046). **j**, Schematic of experimental setup for electroporation of CaMKIIα+ pyramidal neurons with fluorescently tagged (Alexa Fluor 488) anionic dextran. Inset: widefield fluorescence image of Voltron2608-ST fluorescence of targeted neuron; scale bar: 10 µm. **k**, Successful electroporation of neurons with anionic dextran verified by observing fluorescence restricted to the cell of interest; scale bars: 10 µm. **l**, Activation of endogenous GABAARs during Voltron2608-ST voltage imaging was used to estimate DFGABAA from the pyramidal neuron in ‘j’ and ‘k’, both pre-electroporation (pre) and post-electroporation (post). **m**, Population data showing that addition of impermeant anions had no effect on estimated DFGABAA (paired t-test, *P* = 0.2506, *n* = 12). ns = not significant (*P* > 0.05); \**P* ≤ 0.05; \*\*\**P* ≤ 0.001; \*\*\*\**P* ≤ 0.0001; error bars indicate mean ± SEM.

To empirically test these predictions, we used ORCHID strategies to measure the effect of manipulations upon DFGABAA in hippocampal pyramidal neurons. First, we observed that the addition of the selective inhibitor of KCC2, VU0463271 (VU; 10 μM), caused a robust depolarizing shift in estimated DFGABAA (**Fig. 3f-i**). This was evident when DFGABAA was measured either via activation of GABAARs or all-optically, and the effect upon DFGABAA reversed following washout of VU (**Fig. 3f-i**). Second, we assessed DFGABAA before and after the addition of intracellular impermeant anions, which was achieved by single-cell electroporation of fluorescently tagged anionic dextrans (Alexa Fluor 488, see **Methods**) (**Fig. 3j**). Successful addition of impermeant anions was confirmed by directly observing the fluorescent dextrans within the neuron of interest, and DFGABAA was estimated from the size and polarity of GABAAR responses before and after the electroporation procedure (**Fig. 3k-l**). Consistent with the prediction from the modelling work, the addition of impermeant anions had no detectable effect upon the experimentally determined DFGABAA (**Fig. 3m**). Therefore, ORCHID strategies are able to confirm theoretical predictions about the mechanisms that establish DFGABAA.

### ORCHID reveals dynamic ion driving forces over different timescales

In neurons, [Cl^-^]i and EGABAA are considered dynamic variables under the control of multiple activity-dependent processes, and are thought to vary over a range of timescales^4^. We therefore sought to use ORCHID to investigate DFGABAA dynamics, and first examined periods of elevated network activity, during which intense activation of GABAARs is hypothesized to cause transient changes in DFGABAA5. ORCHID was expressed in CaMKIIα+ pyramidal neurons or GAD2+ interneurons, and we used low-Mg^2+^ artificial cerebrospinal fluid (aCSF) to induce seizure-like events (SLEs) in mouse organotypic hippocampal brain slices. DFGABAA was estimated using ORCHID in target cells, whilst a simultaneous whole-cell current-clamp recording from a nearby CA1/CA3 pyramidal neuron provided an independent readout of network activity (**Fig. 4a-d**). The SLEs were characterized by pronounced depolarizing shifts in Vm and intense action potential firing, which could be observed both in the current-clamp recordings and in ORCHID’s voltage signals (**Fig. 4a,c**). CaMKIIα+ pyramidal neurons showed substantial activity-dependent changes in DFGABAA, exhibiting shifts in polarity from a hyperpolarizing DFGABAA at baseline, to a depolarizing DFGABAA immediately post-SLE, which then returned to a hyperpolarizing DFGABAA > 2 min after the SLE (**Fig. 4b**). Similarly, GAD2+ interneurons exhibited a hyperpolarizing DFGABAA at baseline that flipped polarity to become a depolarizing DFGABAA immediately post-SLE, and subsequently returned to a hyperpolarizing DFGABAA after > 2 min (**Fig. 4d**). These dynamics were confirmed by using endogenous GABAAR activation to estimate DFGABAA in CaMKIIα+ pyramidal neurons and GAD2+ interneurons (**Supplementary Fig. 6**). Furthermore, shifts in DFGABAA have been postulated to occur immediately prior to seizure onset and to form part of the mechanism by which seizures initiate^33^. In line with this, ORCHID detected more modest decreases in the magnitude of DFGABAA between baseline and immediately prior to the SLE (< 5 s prior to SLE onset) in both CaMKIIα+ pyramidal neurons and GAD2+ interneurons (**Fig. 4b,d**), suggesting that DFGABAA begins to break down before SLE onset.

**Figure 4:**
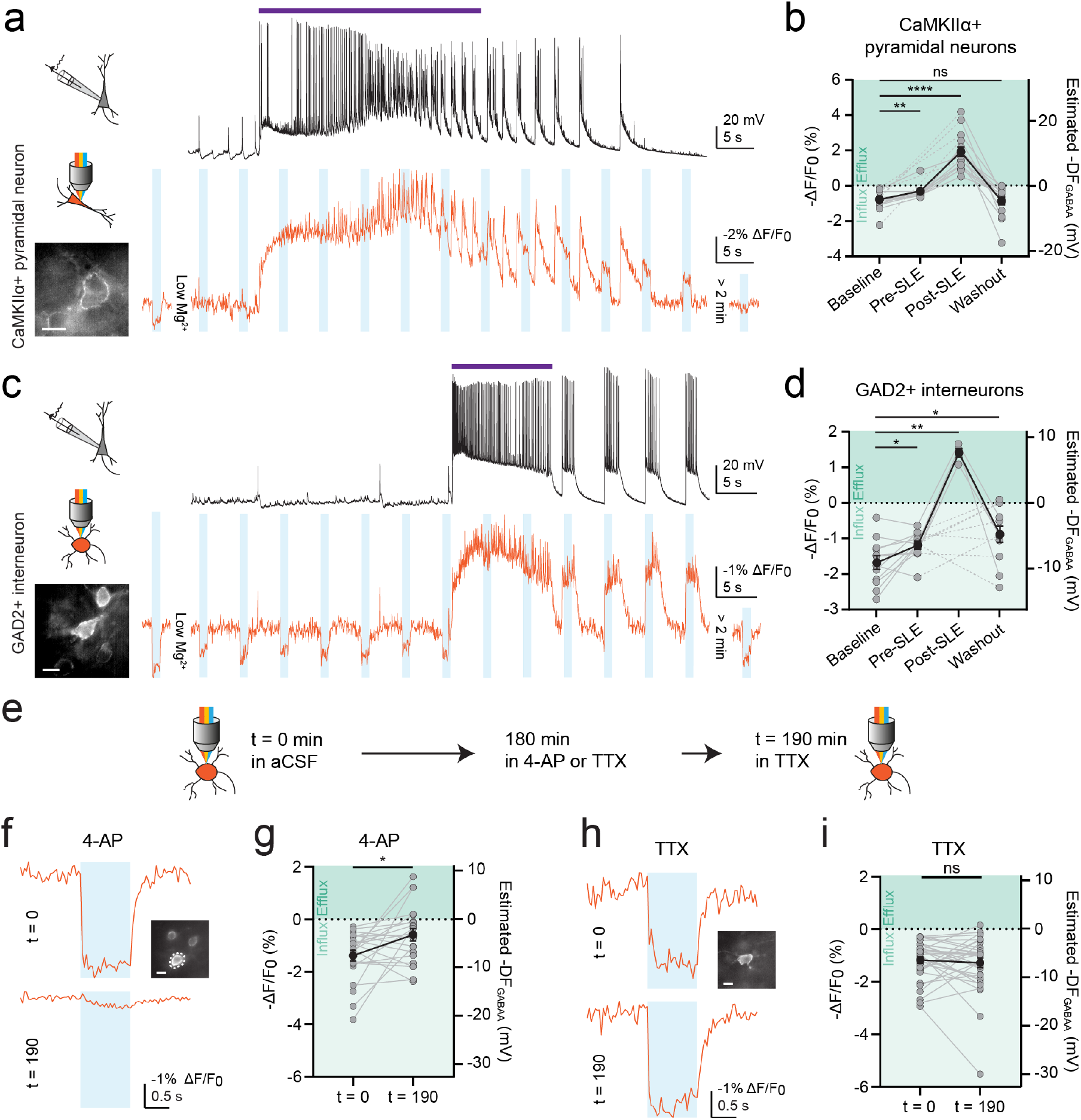
ORCHID reveals dynamic inhibitory receptor driving forces over different timescales. **a**, ORCHID used to investigate activity-dependent variation in DFGABAA in a CaMKIIα+ pyramidal neuron (inset; scale bar: 10 µm, bottom; orange trace) during low-Mg^2+^ induced SLEs. Whole-cell current-clamp recording from a different CA1/3 pyramidal neuron provided an independent readout of SLEs (top; black trace, purple bar indicates SLE). **b**, Population data from CaMKIIα+ pyramidal neurons showing a significant shift in DFGABAA from baseline as compared to immediately pre-SLE (< 5 s before onset, Wilcoxon matched-pairs signed rank test, *P* = 0.0020, *n* = 10) and immediately post-SLE (< 20 s after SLE cessation, Wilcoxon matched-pairs signed rank test, *P* < 0.0001, *n* = 15). However, > 2 min post-SLE DFGABAA had returned to baseline levels (baseline vs washout: Wilcoxon matched-pairs signed rank test, *P* = 0.5995, *n* = 15). Dotted lines indicate cells for which pre-SLE DFGABAA was not recorded (*n* = 5/15). **c**, As in ‘a’ but with GAD2+ interneurons targeted for ORCHID recording. **d**, Population data of DFGABAA in GAD2+ interneurons showed a significant depolarizing shift in DFGABAA between the baseline recordings and the recordings made pre-SLE (paired t-test, *P* = 0.0234, *n* = 12). In 8/12 cells, stimulation of GtACR2 as part of ORCHID caused slice-wide burst firing after the occurrence of a SLE, which persisted for minutes. In the 4/12 recordings where GtACR2-induced burst firing did not occur, there was a significant depolarizing shift in DFGABAA post-SLE (paired t-test, *P* = 0.0047, *n* = 4); dotted lines indicate cells in which burst-firing occurred and post-SLE DFGABAA could not be recorded (*n* = 8/12). When recorded again > 2 min after the SLE, DFGABAA had returned to near baseline values (baseline vs washout, paired t-test, *P* = 0.0317, *n* = 12). **e**, Schematic of experimental design with ORCHID used to make paired recordings of DFGABAA 190 min apart in the same GAD2+ interneurons in organotypic hippocampal brain slices. **f**, Top, baseline recording before a slice was incubated in 4-AP (50 µM) for 180 min. Bottom, recording after the slice had been transferred to aCSF containing TTX (1 µm) for 10 min. Inset: widefield fluorescence image of recorded neuron; scale bar: 10 µm. **g**, Population data showing DFGABAA depolarized significantly after a 180 min incubation in 4-AP (Wilcoxon matched-pairs signed rank test, *P* = 0.0229, *n* = 22). **h**, As in ‘f’ but with recordings of DFGABAA before and after a 190 min incubation in TTX. Inset: widefield fluorescence image of recorded neuron; scale bar: 10 µm. **i**, Population data showing magnitude of DFGABAA was not significantly different after a 190 min incubation in TTX (Wilcoxon matched-pairs signed rank test, *P* = 0.5888, *n* = 34). ns = not significant (*P* > 0.05); \**P* ≤ 0.05; error bars indicate mean ± SEM.

Having demonstrated ORCHID’s ability to report DFGABAA dynamics on a timescale of seconds, we next sought to assess ORCHID’s potential to measure longer-term changes in a neuron’s DFGABAA, such as those that have been linked to activity-dependent changes in CCCs^34^. To this end, ORCHID was used to track DFGABAA in the same GAD2+ interneurons before and after a 180 min treatment with either 4-aminopyridine (4-AP; 50 µM) to increase neuronal activity or tetrodotoxin (TTX; 1 µM) to silence neuronal activity (**Fig. 4e**). When DFGABAA measurements were made after the 180 min treatment and a subsequent 10 min incubation in TTX, ORCHID revealed that raising neuronal activity had caused a sustained and robust depolarizing shift in neuronal DFGABAA (**Fig. 4f,g**), which was not evident in neurons that had been silenced with TTX (**Fig. 4h,i**), or in neurons exposed to vehicle control for the equivalent 180 min period (**Supplementary Fig. 7**). Hence, ORCHID can provide high-throughput measurements of resting and dynamic DFGABAA across timescales of seconds and hours.

### Astrocytes sustain an outward anion driving force during enhanced network activity

Astrocytes are known to play important roles in regulating and maintaining neuronal ion gradients by modulating extracellular ion concentrations. However, due to the difficulty in performing electrophysiological recordings from this cell type, it has not previously been possible to measure astrocytic DFGABAA across different types of network activity. Using the Cre-lox system, we restricted ORCHID expression to GFAP+ astrocytes in mouse organotypic hippocampal brain slices and optically estimated DFGABAA. We observed that, in stark contrast to neurons, the astrocytic DFGABAA under baseline conditions was strongly depolarizing (**Fig. 5a-c**). A related idea that has been proposed is that astrocytes may serve to continuously supplement anions into the extracellular space during periods of enhanced network activity, and thereby help to maintain the effectiveness of GABAAR-mediated inhibitory synaptic transmission^14^. To test this explicitly, we used ORCHID to continuously monitor astrocytic DFGABAA across periods of intense network activity, which was achieved by inducing SLEs through low-Mg^2+^ aCSF. Whilst there was a modest decrease in the magnitude of DFGABAA during the SLE, ORCHID revealed that the astrocytes continued to exhibit a strongly depolarizing DFGABAA before and after the SLE, and are therefore capable of maintaining a driving force for anion efflux during substantial changes in network activity (**Fig. 5d,e**). When compared to neurons, such markedly different resting and dynamic DFGABAA in astrocytes suggests that they differ in how they utilize major Cl^-^ regulatory mechanisms such as NKCC1 and KCC2, which act to transport Cl^-^ in and out of the cell respectively. To assess this, we performed single-nucleus RNA sequencing (snRNAseq) on the same hippocampal brain slice preparations (see **Methods**) and distinguished pyramidal neurons and astrocytes using unbiased clustering of the transcriptomic profiles (**Fig. 5f**). After normalizing for total gene expression, we observed that KCC2 (Slc12a5) was more highly expressed in pyramidal neurons than astrocytes. Meanwhile NKCC1 (Slc12a2), although comparatively low in both cell types, exhibited higher expression in astrocytes than pyramidal neurons (**Fig. 5f**). Thus, ORCHID reveals unique features of astrocytic DFGABAA compared to neurons, that correlate with differences in their Cl^-^-transport mechanisms.

**Figure 5:**
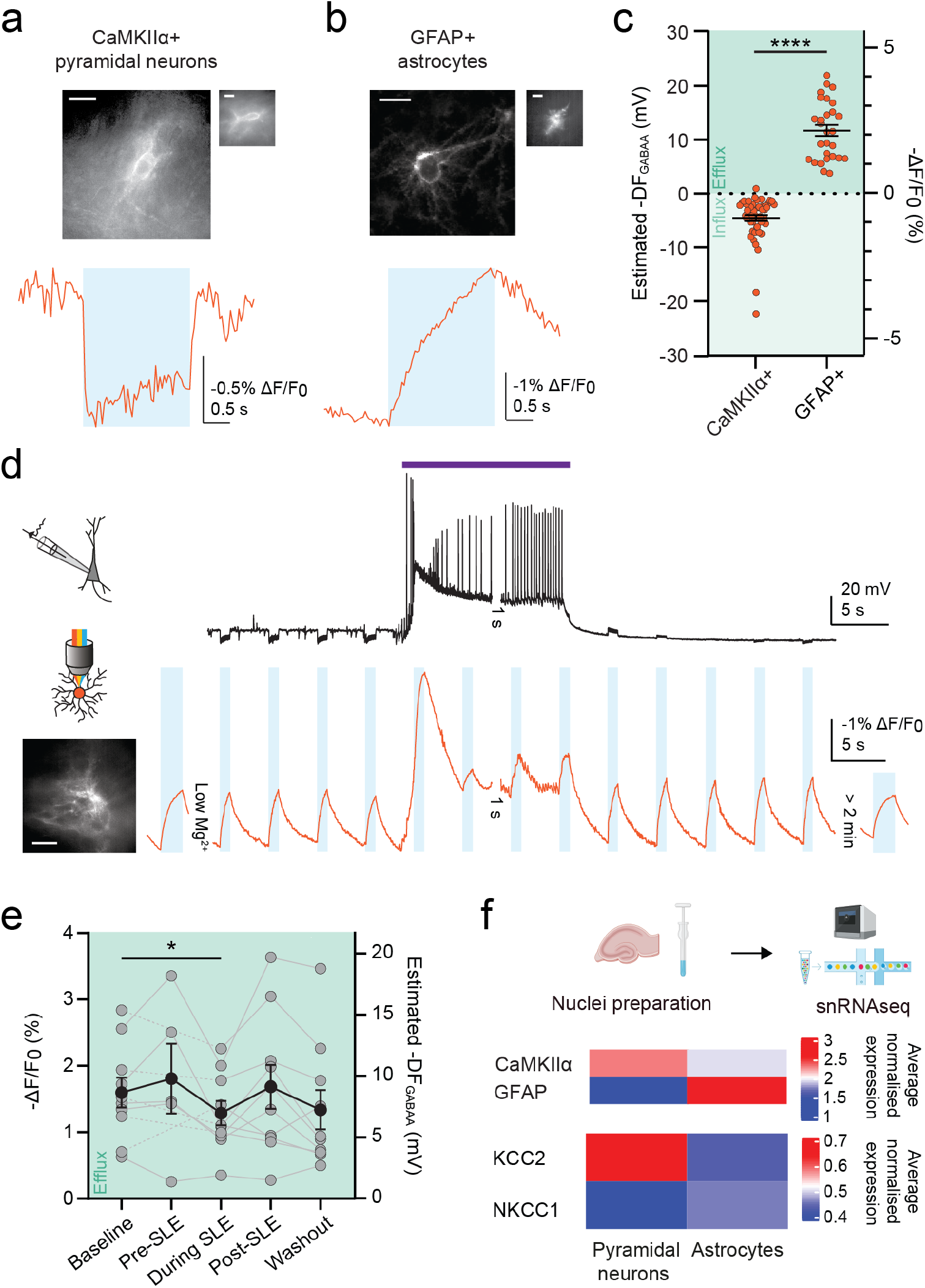
Astrocytes sustain an outward anion driving force during enhanced network activity. ORCHID was used to compare DFGABAA in CaMKIIα+ pyramidal neurons (**a**) and GFAP+ astrocytes (**b**). Widefield and confocal images of cells expressing ORCHID (top) and fluorescence recordings used to estimate DFGABAA from each cell type (bottom), with the cell that was recorded from shown inset; scale bar: 10 µm. **c**, Estimated DFGABAA was significantly different between GFAP+ astrocytes and CaMKIIα+ pyramidal neurons (-11.73 ± 1.025 mV vs 4.52 ± 0.57 mV; Mann-Whitney test, *P* < 0.0001, *n* = 28 and 51 cells respectively). **d**, ORCHID was used to investigate network activity-dependent variation in DFGABAA in GFAP+ astrocytes during low-Mg^2+^ induced SLEs (inset; scale bar: 10 µm, bottom; orange trace). A whole-cell current-clamp recording from a CA1/3 pyramidal neuron provided a readout of network activity (top; black trace, purple bar indicates SLE). **e**, Population data of DFGABAA in GFAP+ astrocytes showed depolarizing DFGABAA at baseline that were maintained pre-SLE (paired t-test, *P* = 0.3098, *n* = 5), post-SLE (paired t-test, *P* = 0.5813, *n* = 10), and after washout (Wilcoxon matched-pairs signed rank test, *P* = 0.0840, *n* = 10). However, there was a small decrease in the magnitude of DFGABAA during the SLE (baseline vs during SLE: paired t-test, *P* = 0.0334, *n* = 10). Dotted lines indicate cells for which pre-SLE DFGABAA was not recorded (*n* = 5/10). **f**, Top, schematic showing nuclear dissociation and snRNAseq from mouse organotypic hippocampal brain slices performed using the 10X Genomics platform. Middle, heatmap showing the average normalized expression of CaMKIIα and GFAP RNA in the pyramidal neuron and astrocyte clusters respectively. Bottom, heatmap showing the average normalized gene expression between pyramidal neurons vs astrocytes, which was different for KCC2 (Slc12a5, *P =* 0.0324*, n =* 4 samples) and NKCC1 (Slc12a2, *P =* 0.0087*, n =* 4 samples). ns = not significant (*P* > 0.05); \**P* ≤ 0.05; \*\*\*\**P* ≤ 0.0001; error bars indicate mean ± SEM.

### *In vivo* determination of resting and dynamic inhibitory receptor driving forces in neurons

Finally, ORCHID offers the opportunity to generate estimates of undisturbed DFGABAA in the intact brain. We therefore assessed ORCHID’s ability to report resting and dynamic DFGABAA *in vivo*. ORCHID was expressed in layer 1 (L1) GAD2+ interneurons in the mouse primary somatosensory cortex (S1), and anesthetized *in vivo* imaging was combined with simultaneous local-field potential (LFP) recordings to provide an independent measure of local population activity (**Fig. 6a**; **Methods**). Under baseline conditions, we observed a range of DFGABAA polarities and magnitudes across different neurons, including hyperpolarizing, shunting, and depolarizing DFGABAA (**Fig. 6b,c**). In a subset of mice, we performed continuous DFGABAA measurements before and after infusing 4-AP (500 µM) into the cortex to elicit SLEs (**Methods**). The occurrence of SLEs was confirmed from the LFP recordings and DFGABAA could be tracked in the same individual neurons before and after the SLE (**Fig. 6d**). This revealed that L1 GAD2+ interneurons exhibit depolarizing shifts in DFGABAA as a result of SLEs (**Fig. 6e**). These data demonstrate that *in vivo* ORCHID affords easy and rapid reporting of resting and activity-dependent dynamic shifts in DFGABAA in the intact brain, under control and disease-relevant conditions.

**Figure 6:**
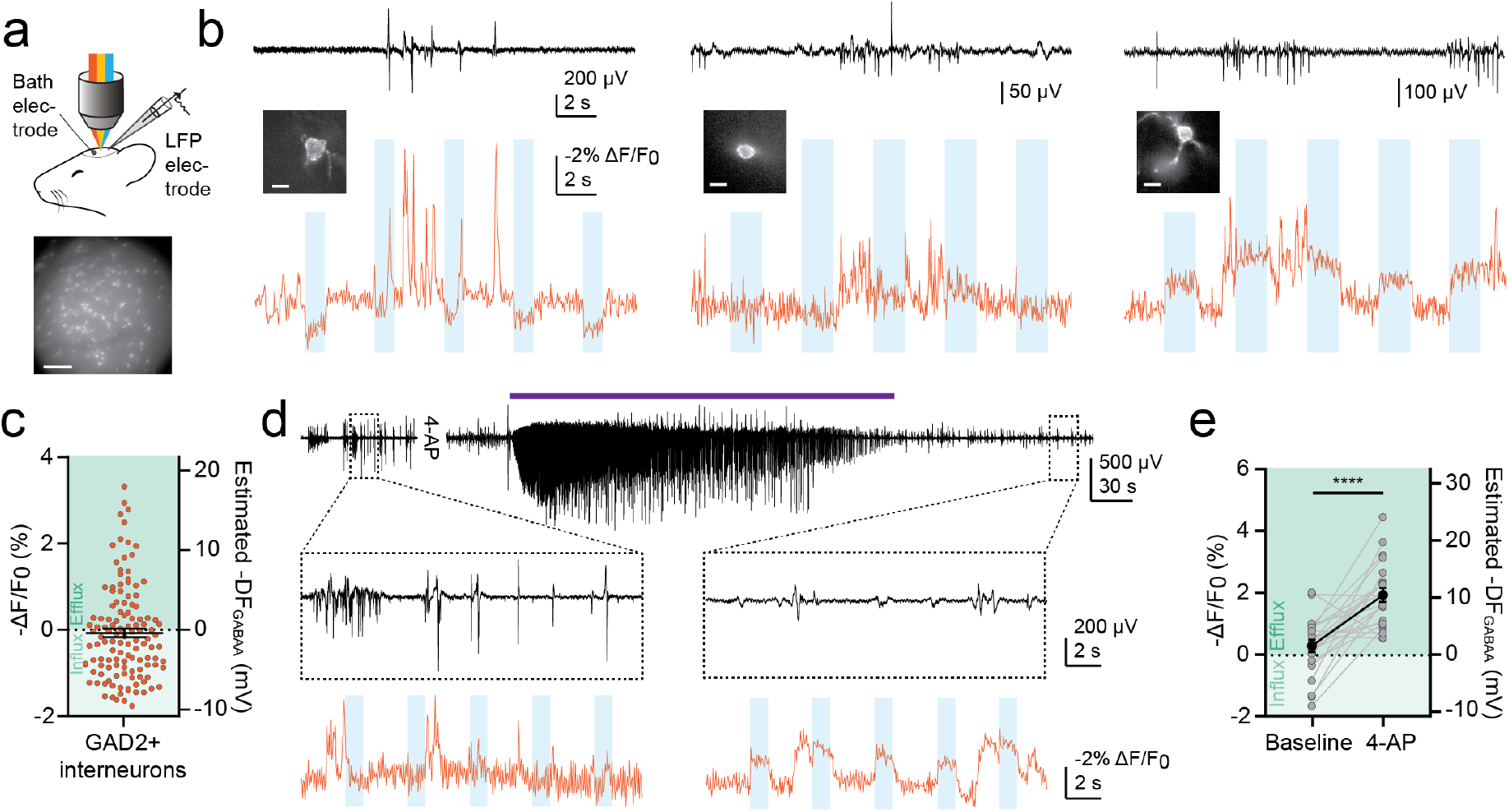
In vivo determination of resting and dynamic inhibitory receptor driving forces in neurons. **a**, Schematic of experimental setup (top) and a widefield fluorescence image of a region of L1 in S1 with GAD2+ interneurons expressing ORCHID (bottom); scale bar: 100 µm. **b**, LFP recordings were used to independently record population activity (black traces; top). ORCHID was used to concurrently record DFGABAA in GAD2+ interneurons in L1 of the anesthetized mouse brain (orange traces; bottom) where a variety of different DFGABAA values were recorded including hyperpolarizing (left), shunting (center) and depolarizing DFGABAA (right). Inset: widefield fluorescence images of the cells from which recordings were made; scale bar: 10 µm. **c**, Population data showing DFGABAA in L1 interneurons in S1 (0.3873 mV ± 0.5348 mV, *n* = 123 cells from 10 mice). **d**, 4-AP (500 µM) was infused into the cortex buffer in the recording chamber over the craniotomy to investigate DFGABAA dynamics during SLEs *in vivo*. ORCHID recording from a GAD2+ interneuron before (left) and after (right) a 4-AP induced SLE (bottom; orange traces). A LFP recording provided readout of network activity (black traces, purple bar indicates SLE). **e**, Population data of DFGABAA in GAD2+ interneurons showed a large depolarizing shift in DFGABAA post-SLE (paired t-test, *P* < 0.0001, *n* = 22 cells from 4 mice). \*\*\*\**P* ≤ 0.0001; error bars indicate mean ± SEM.

## Discussion

All-optical approaches that combine voltage imaging and optogenetic stimulation, represent a rapidly emerging strategy for studying neuronal dynamics in the intact nervous system^35,36^. Here we present the first all-optical strategy for studying ionic driving forces. We describe ORCHID - a powerful all-optical tool for the estimation of DFGABAA, which represents the electromotive force for ion flux across inhibitory receptors. Whilst ORCHID could be used to estimate DFGABAA in any cell type or organ, we validate its use in the rodent nervous system. Taking advantage of ORCHID’s ability to provide high-throughput, genetically targeted estimates of DFGABAA over extended periods of time, we confirm theoretical predictions regarding the mechanisms that establish DFGABAA, utilize ORCHID to reveal novel, cell-type-specific differences in activity-dependent DFGABAA changes during network activity, and provide the first *in vivo* measurements of intact DFGABAA in mouse cortical neurons.

ORCHID is easy to use with no spectral crosstalk between the orange-light excitation of the GEVI, Voltron2608-ST, and blue-light activation of the integrated anion-permeable channel, GtACR2. This means that ORCHID is compatible with a wide variety of experimental paradigms. ORCHID’s accuracy in reporting DFGABAA depends upon the amplitude of the light-activated anion conductance, relative to other conductances that are active in the cell of interest. For this reason it is important that GtACR2 is known to generate large transmembrane conductances^24^, and any future optimization of light-activated anion channels are predicted to further improve ORCHID’s accuracy. GtACR2’s permeability sequence to different anions (NO3-> I^-^ > Br^-^ > Cl^-^ > F^−^) is the same as other anion channels including GABAARs and GlyRs^24–26^, which are also permeable HCO3-, and exhibit a Cl^-^ to HCO3-permeability ratio of five to one ^25,26,37,38^. The equivalence of the anion permeabilities of GtACR2’s and GABAARs is supported by the fact that their reversal potentials are the same^27,28^, and also our evidence that ORCHID measurements of DFGABAA are indistinguishable to those made by activating GABAARs. ORCHID therefore provides an effective readout of the driving force across the receptors that mediate fast synaptic inhibition in the brain. As GtACR2 is primarily permeable to Cl^-24^, ORCHID measurements can also be considered to be estimates of Cl^-^ driving force (DFCl), and future efforts to modulate the ion selectivity of light-activated Cl^-^ channels could further enhance the accuracy of all-optical reporters of DFCl.

ORCHID’s all-optical approach has several powerful advantages over currently available techniques for estimating DFGABAA. Firstly, and most fundamentally, ORCHID does not perturb ionic gradients prior to measurement. Secondly, it is both genetically targetable and addressable by light, which affords excellent temporal and spatial control. Thirdly, ORCHID’s high-throughput nature allows for measurements that are orders-of-magnitude faster than traditional techniques. A typical gramicidin-perforated patch-clamp recording requires 30-60 minutes per cell, compared to 1-5 seconds for an ORCHID recording. And fourthly, ORCHID affords measurements from small cells and potentially subcellular compartments, which are difficult to access using electrode-based techniques. In theory, a fluorescent reporter of [Cl^-^]i could be used to report relative changes in a cell’s DFGABAA, by measuring changes in the amplitude of the [Cl^-^]i response to an evoked Cl^-^ conductance^39^. However, this is unlikely to be achieved in practice because of the inherent limitations in the sensitivity of Cl^-^ imaging^22^. Furthermore, detecting [Cl^-^]i changes would require much larger and longer conductances that alter [Cl^-^]i in the cytoplasm, unlike ORCHID which only reflects Cl^-^ movement across the membrane. Finally, a fluorescent reporter of [Cl^-^]i would not account for the HCO3-component of DFGABAA. By integrating a transmembrane voltage-reporter element and an anion conductance, ORCHID is highly sensitive to transmembrane flux and able to simultaneously readout Vm, which are key to making accurate measurements of DFGABAA.

Our results demonstrate how ORCHID strategies can be harnessed to test theoretical predictions about the underlying mechanisms that establish ionic driving forces. Our computational modelling and empirical evidence support the view that DFGABAA is primarily established by the action of CCCs such as KCC2^1^. Therefore, whilst genetically encoded Cl^-^ indicators and other techniques that directly report [Cl^-^]i are of considerable value, ORCHID highlights the danger of interpreting [Cl^-^]i changes in the context of inhibitory synaptic transmission and DFGABAA, without a concurrent readout of Vm. This is particularly the case for manipulations such as altering impermeant anions, which are predicted to shift both ECl, EGABAA and Vm in tandem^32,40^. Given its ability to provide a sensitive readout of DFGABAA, we note that ORCHID could serve as a high-throughput means for discovering new and improved enhancers, or inhibitors, of transmembrane Cl^-^ transport.

By avoiding intracellular recordings, ORCHID enabled us to monitor unperturbed resting and dynamic DFGABAA. We confirm predictions from previous work by showing that intense network activity can induce a profound change in DFGABAA polarity in neurons, with GABAergic synaptic input switching from inhibitory to excitatory, which is relevant for the management of status epilepticus^41,42^. Whilst SLE-associated DFGABAA estimates have been made using gramicidin-perforation in pyramidal neurons^15^, we provide the first data showing that inhibitory interneurons also undergo an inversion in DFGABAA following seizure-like activity. Recent work has suggested that the transition to seizures is associated with incremental increases in neuronal [Cl^-^]i, which is predicted to reduce DFGABAA prior to seizure onset^33^. We provide direct evidence for this clinically relevant phenomenon, and thereby demonstrate the utility of ORCHID in dynamically estimating DFGABAA across network states.

By harnessing ORCHID’s ability to be genetically targeted, we recorded DFGABAA from multiple cell types including large populations of hippocampal CaMKIIα+ pyramidal neurons, GAD2+ interneurons, and GFAP+ astrocytes. This revealed that hippocampal astrocytes display a strong depolarizing DFGABAA at rest, which is opposite to the hyperpolarizing DFGABAA exhibited by hippocampal pyramidal neurons *in vitro*, and is supported by the differential expression of CCCs between these two cell populations^15,27,43^. It has been suggested that astrocytes could serve as a source of extracellular anions that help to sustain the effectiveness of inhibitory GABAergic transmission^44^, which has recently been supported by direct measurements of astrocytic [Cl^-^]i during periods of network activity^45^. In support of this idea, our measurements of dynamic DFGABAA reveal for the first time that astrocytes are able maintain a force for the outward flux of anions, even during intense network activity. As other cell types exhibit specific expression patterns of CCCs and other anion transporters^46^, ORCHID could be used to determine how these translate into differences in the anion driving forces in cells such as oligodendrocytes, ependymal cells, microglia, and endothelial cells. Finally, we also demonstrate ORCHID’s ability to provide measurements of undisturbed DFGABAA in the intact brain. Our *in vivo* results captured a range of resting DFGABAA values in cortical L1 GAD2+ interneurons and revealed that DFGABAA is a dynamic parameter in the intact brain, with periods of intense activity causing substantial shifts towards depolarizing DFGABAA. Indeed, through the use of fiber photometry, or the implementation of a GEVI that is suitable for use with 2-photon microscopy, ORCHID has the potential to generate measurements of resting and dynamic DFGABAA throughout the mammalian brain^36^, including from deep brain structures.

In terms of future applications, a further important question is how DFGABAA is regulated at a subcellular level. For example, it has been suggested that the properties of inhibitory synaptic input vary across a neuron and between different cellular compartments^15,47–50^. However, this has been difficult to address given the issue of space-clamp and additional technical challenges associated with electrode-based recordings. One could imagine combining ORCHID strategies with protein-targeting motifs and the patterned delivery of light in order to record DFGABAA from difficult-to-access subcellular compartments, such as distal apical dendrites, the axon initial segment, and astrocytic processes. In a similar manner, one could adopt targeted ORCHID strategies to estimate DFGABAA / DFCl across the membranes of organelles, such as mitochondria, lysosomes and vesicles. In doing so, ORCHID could be used to study a range of cellular phenomena, including cell division, growth and migration, in which anion / Cl^-^ fluxes across different membranes contribute to the underlying intracellular and intercellular signaling processes.

In conclusion, we establish ORCHID as an all-optical strategy for estimating DFGABAA in the nervous system. This underscores the potential for similar all-optical strategies for elucidating how ionic driving forces affect cellular signaling in a range of biological contexts.

## Methods

### ORCHID construct subcloning

The ORCHID construct combined GtACR2 from the pAAV-EF1α-FRT-FLEX-GtACR2-EYFP plasmid, which was a gift from Mingshan Xue (Addgene plasmid #114369) (Messier *et al*., 2018), with Voltron2-ST from the pGP-pcDNA3.1 Puro-CAG-Voltron2-ST plasmid (Abdelfattah *et al*., 2023), which was kindly provided by Eric Schreiter and Ahmed Abdelfattah. The two genes were linked by a P2A linker sequence identical to that used by Holst *et al*. (2006) to form an insert sequence, and include the Kozak sequence 5’ GCCACC(ATG) 3’. The vector used for the ORCHID construct originated from pAAV-hSyn-DIO {ChETA-mRuby2}on-W3SL, which was a gift from Adam Kepecs (Addgene plasmid #111389) (Li *et al*., 2018), with the hSyn promoter replaced with the EF-1α promoter from pAAV-nEF Con/Foff hChR2(H134R)-EYFP, which was a gift from Karl Deisseroth (Addgene plasmid #55647) (Fenno *et al*., 2014). The GtACR2-P2A-Voltron2-ST sequence of the ORCHID construct was synthesized in antisense (Thermo Fisher Scientific GeneArt Gene Synthesis) and inserted into the vector using the Bsp1407I/NheI restriction enzyme sites and the FastDigest versions of the two enzymes (Thermo Fisher Scientific). Sanger sequencing confirmed the sequence of the final construct. Production of the AAV vector (serotype 1) containing the completed pAAV-nEF-DIO-GtACR2-P2A-Voltron2ST-W3SL construct was carried out by Dr R. Jude Samulski and the University of North Carolina Vector Core.

### Mouse hippocampal organotypic brain slice cultures

The animals used in this study were wild-type (C57BL/6 background) or GAD2-IRES-Cre mice (RRID:IMSR_JAX:010802), with the GAD2-IRES-Cre strain being characterized by expression of Cre recombinase in all GABAergic interneurons. The use of these animals was approved by the University of Cape Town Animal Ethics Committee (AEC Protocol 021/026 and AEC Protocol 022/038). Organotypic brain slice cultures were prepared using postnatal day (P) 7 mice and followed the protocol originally described by Stoppini, Buchs and Muller (1991) [for details see Raimondo *et al*. (2016)] All reagents were purchased from Sigma-Aldrich unless otherwise stated. Briefly, the mice were killed, and the brains were removed and placed in 4°C Earle’s Balanced Salt Solution (EBSS), with 6.1 g/l HEPES, 6.6 g/l glucose, and 5% 1 M NaOH. The hippocampi were removed and sectioned into 350 μm slices using a tissue chopper (McIlwain). The slices were placed on Millicell-CM membranes and cultured in medium consisting of (v/v): 25% EBSS; 49% minimum essential medium; 25% heat-inactivated horse serum; 1% B27 (Invitrogen, Life Technologies), and 6.2 g/l D-glucose. The slices were incubated at 35-37°C in a 5% CO2 humidified incubator.

Viral transfection was performed at 0-1 days *in vitro* (DIV). For experiments in which endogenous GABAARs were activated, pGP-AAV-syn-flex-Voltron2-ST-WPRE-SV40 was utilized, while for ORCHID experiments pAAV-nEF-DIO-GtACR2-P2A-Voltron2ST-W3SL was utilized. AAV-CaMKIIα-Cre-GFP and AAV8-GFAP-GFP-Cre were used to target CaMKIIα+ pyramidal neurons and GFAP+ astrocytes respectively. Here, as in all relevant methods, micropipettes were prepared from borosilicate glass capillaries (Warner Instruments). The AAVs were loaded into a glass micropipette along with FastGreen (0.1% w/v) to aid visualization, and were injected into the slices using the Openspritzer system (Forman *et al*., 2017). Biolistic transfection was performed at 4-5 DIV. Cartridges were prepared as described by O’Brien and Lummis (2006), using 20 mg of pGP-pcDNA3.1 Puro-CAG-Voltron2-ST plasmid DNA and 1.6 μm gold microcarriers (Bio-Rad). Slices were transfected using a 3D-printed version of the modified barrel design (O’brien *et al*., 2001).

Recordings were performed at 7-14 DIV, which is equivalent to P14-21. This, as well as previous work, has shown that pyramidal neurons in the organotypic slices of the hippocampus have mature and stable Cl^-^ homeostasis mechanisms at this stage, as evidenced by their hyperpolarizing DFGABAA (Streit, Thompson and Gähwiler, 1989; Ilie, Raimondo and Akerman, 2012; Raimondo *et al*., 2012; Wright *et al*., 2017). Prior to imaging, slices were incubated for 30 min in aCSF bubbled with carbogen gas (95% O2: 5% CO2), containing either JF549 or JFX608 dye at 1 mM. The aCSF was composed of (in mM): NaCl (120); KCl (3); MgCl2 (2); CaCl2 (2); NaH2PO4 (1.2); NaHCO3 (23); and D-glucose (11). During this incubation step the dye binds irreversibly to the Voltron2-ST protein (Abdelfattah *et al*., 2019). After the incubation was complete, slices were transferred to a submerged recording chamber where they were continuously superfused with 32°C carbogen-bubbled aCSF using a peristaltic pump (Watson-Marlow). Pharmacological agents included CGP-55845 (5 µM) and 2-chloroadenosine (4 µM) in the recording aCSF for all experiments in which GABAARs were activated, as well as pictrotoxin (PTX; 100 µM), VU0463271 (VU; 10 μM), 4-aminopyradine (4-AP, 50 µM), and tetrodotoxin (TTX; 1 µM) for subsets of the *in vitro* experiments.

### Surgical procedures for in vivo imaging

Intracranial bulk regional viral injections of pAAV-nEF-DIO-GtACR2-P2A-Voltron2-ST-W3SL were performed on P0-1 GAD2-IRES-Cre mouse pups (Cheetham, Grier and Belluscio, 2015; Oomoto *et al*., 2021). Adult breeders were removed from the home cage and placed in a holding cage for the duration of the procedure. A 30G insulin syringe was loaded with 1 μl AAV combined with FastGreen (0.1% w/v) to aid visualization and mounted in a stereotaxic frame. Pups were individually removed from their home cage and anesthetized using 2 ml isoflurane pipetted onto a gauze swab in an enclosed space. Respiration rate and skin color were monitored to assess depth of anesthesia. Once depth of anesthesia was sufficient, pups were rapidly transferred to a heating pad beneath the stereotaxic frame and the AAV was injected into S1 of the left cerebral hemisphere at a depth of 1-2 mm, with a successful injection indicated by green coloration of the hemisphere. The injection typically took ∼30 s. The pup was then transferred back to its home cage and monitored until fully recovered from anesthesia, when the injection of the next pup was begun. We observed a 100% survival rate of injected pups using this method.

Imaging experiments began after P28. Retroorbital injections of JF608 were performed 3-24 h prior to imaging (Abdelfattah *et al*., 2019, 2023). 100 nmol JF608 was dissolved in 20 μl DMSO, 20 μl Pluronic F-127 (20% w/v in DMSO, Thermo Fisher Scientific), and 60 μl sterile phosphate-buffered saline (PBS). The mice were anesthetized using an intraperitoneal (IP) injection of ketamine/xylazine mixture (ketamine 80 mg/kg and xylazine 10 mg/kg). The JF608 solution was delivered using a 30G insulin syringe as described by Yardeni *et al*. (2011). The preparation for anesthetized recordings was adapted from Burman *et al*. (2023). Mice were anesthetized using 3 l/s O2 with 5% isoflurane for induction, and with the isoflurane reduced to 1-2% for maintenance. Once anesthetized, the animal was secured in a stereotaxic frame (Narishige). An IP injection of buprenorphine (0.5 mg/kg) and a subcutaneous (SC) injection of 200 μl sterile saline were administered for analgesia and hydration respectively. The mouse’s body temperature was maintained at ∼37°C using a heating pad and a temperature probe. The head was shaved, and eye-protecting ointment (Duratears) was applied to each eye. An incision in the scalp was made using surgical scissors and the area was expanded by blunt dissection to expose the skull. Forceps were used to remove any membranous tissue from the surface of the skull. Tissue adhesive (Vetbond) was used to fix the edges of the incised scalp to the skull and to stabilize the cranial sutures. Several layers of dental cement (InterDent) were applied to create a recording chamber on top of the skull. A 0.5 mm craniotomy was drilled over S1 using a dental drill (Dental Lab). The craniotomy was submerged in cortex buffer containing (in mM): NaCl (125), KCl (5), HEPES (10), MgSO4·7H2O (2), CaCl2·2H2O (2), D-glucose (10). The bone flap and dura were removed. The mouse was then transferred to the imaging setup. In a subset of animals, 4-AP was infused into the cortex buffer in the recording chamber using a pipette (500 µM working concentration). The recording session typically lasted 3 h.

### Electrophysiology

For patch-clamp recordings, micropipettes were filled with a low-Cl^-^ internal solution composed of (in mM): K-gluconate (120); KCl (10); Na2ATP (4); NaGTP (0.3); Na2-phosphocreatinine (10) and HEPES (10). For high-Cl^-^ recordings, pipettes were filled with a high-Cl^-^ internal solution ([Cl^-^] = 141 mM) composed of (in mM): KCl (135), NaCl (8.9) and HEPES (10). Recordings were made using an Axopatch 200B amplifier (Molecular Devices) and data was acquired using WinWCP (University of Strathclyde). For experiments investigating the role of impermeant anions in setting DFGABAA, anionic 10 000 MW dextran bound to Alexa-Flour 488 (Thermo Fisher Scientific) was electroporated into cells. This molecule is a hydrophilic polysaccharide, being both membrane impermeant and having a large negative average charge. A micropipette was filled with 5% dextran solution in PBS, and the micropipette was positioned near the soma of the targeted neuron. Voltage pulses (5-10 pulses, 20 ms duration, 0.5-1 V) were applied using a stimulus isolator, and successful electroporation was confirmed visually by observing the neuron filled with the fluorescent dye. For *in vivo* LFP recordings, micropipettes were filled with cortex buffer. The tip of the pipette was broken against tissue paper with the aim of achieving a tip diameter of ∼5 µm. Recordings were made using a differential amplifier (A-M system, 1800) and data was acquired using LabChart (ADInstruments).

### Widefield fluorescence imaging

Neurons were visualized using a BX51WI upright microscope (Olympus) equipped with a 20x water-immersion objective (Olympus XLUMPlanFL) and a sCMOS camera (Andor Zyla 4.2). For imaging Voltron2549-ST, a green LED with a 555/28 nm excitation filter (Lumencor Aura III) was used for excitation (20.6 mW/mm^2^) with a further multi-band filter set for collecting emission between 590 and 660 nm (Chroma, 69401-ET-380/55-470/30-557/35 multi LED). For Voltron2608-ST or ORCHID, a high-powered orange LED (590 nm, SOLIS-590C, Thorlabs) was used for excitation (143 mW/mm^2^). Optogenetic activation of GtACR2 as a part of ORCHID was achieved using a blue LED (475/28 nm, Lumencor Aura III), with the intensity of this LED fixed at 3.6 mW/mm^2^. A T570lpxr beam splitter was used for combining the blue and orange LEDs, before a filter set with a 599/13 nm excitation filter, a 612 nm long pass beam splitter, and a 632/28 nm emission filter was used (Chroma, 49311-ET-Red#3 Narrow band FISH).

Illumination of the tissue was restricted to a 140 µm diameter region using the built-in Olympus variable-diameter field stop aperture. Image acquisition was controlled using μManager. For current-step recordings, exposure was 5 ms with an image acquisition frequency of 200 Hz. For all voltage-step recordings and Voltron2549-ST recordings, exposure was 10 ms and image acquisition frequency was 100 Hz. For ORCHID recordings exposure was 20 ms and image acquisition frequency was 25 Hz. The imaging region was 256 x 256 pixels, and with no overlap between successive camera exposures this ensured a 20 ms gap between successive exposures. Transistor-transistor logic (TTL) signals from the camera were set to output throughout the exposure, and this signal was sent to an Arduino that ran a custom script to activate the blue LED immediately after each camera exposure ended and for 15 ms, allowing for a 5 ms gap between the blue LED being switched off and the subsequent camera exposure. We found this to be crucial for artefact-free imaging. Camera acquisition and blue-light activation were thus both at a frequency of 25 Hz, with 50% and 37.5% on-time respectively.

When utilizing Voltron2549-ST with GABAAR activation, 500 μM GABA in recording aCSF was loaded into a micropipette and 100 ms puffs of synthetic air were delivered to the micropipette using the Openspritzer system. Imaging traces were 7 s long with a 1 s baseline period prior to the puff. Care was taken to ensure no movement artefact from the puff. ORCHID recordings had a duration of 25 s and consisted of five 1 s or 2 s strobed optogenetic activations, which were averaged post-hoc for some experiments (see Data Analysis and Statistics). During low-Mg^2+^ wash-ins, in order to increase the chance of recording a SLE, the duration of recordings was increased to 65 s, and ORCHID was activated with 1 s periods of strobed light every 5 s. SLEs were defined as events with significant deviation from the resting potential in patch-clamp recordings (> 2 standard deviations) lasting for at least 5 s.

### Confocal imaging

Confocal images of live brain slices were obtained on a LSM 880 Airyscan confocal microscope (Carl Zeiss, ZEN SP 2 software) using a C-Apochromat 40x objective.

### Single-nucleus RNA sequencing

Nuclei from mouse hippocampal organotypic hippocampal brain slices were dissociated in lysis buffer (Nuclei EZ Prep, NUC101) on ice and processed via the 10X Genomics platform to prepare libraries for sequencing as outlined in the Chromium Next GEM 3’ Single Cell 3 Reagent Kits v3.1 user guide. Four samples, each comprising 36 hippocampal slices, were processed. Libraries were sequenced using a Illumina NovaSeq 6000 S2 flow cell. Analysis of the snRNAseq data was performed on facilities provided by the University of Cape Town’s ICTS High Performance Computing team. Cell Ranger version 7.1.0 was used to map paired-end sequencing reads to the mouse reference transcriptome (refdata-gex-mm10-2020-A). The filtered feature-barcode matrix of each sample was converted into a Seurat object. Samples were processed according to the standard Seurat pipeline in R (Hao *et al*., 2021).

Quality control (QC) involved removing nuclei with less than 500 unique molecular identifiers (UMIs), fewer than 250 genes expressed, log10GenesPerUMI less than 0.8, and/or a mitochondrial ratio greater than 0.2. Additionally, all mitochondrial genes were excluded from the dataset. Three doublet identification tools were employed, including DoubletFinder (McGinnis, Murrow and Gartner, 2019), Scrublet (Wolock, Lopez and Klein, 2019), and DoubletDecon (DePasquale *et al*., 2019). For each sample, all doublets called by DoubletFinder together with the intersection of doublets called by Scrublet and DoubletDecon were filtered out. Following QC, normalization and variance stabilization were performed using the SCTransform() function. The mitochondrial ratio was regressed out during this step. Integration of the different samples was performed via Seurat’s FindIntegrationAnchors() function using the top 2000 most variable features and 30 dimensions followed by the IntegrateData() function. A principle component analysis (PCA) was run on the integrated data followed by clustering using the FindNeighbors() and FindClusters() functions.

Several cluster annotation approaches were employed including automated annotation, manual annotation, and Seurat’s label transfer method. For automated annotation, Seurat’s FindAllMarkers() function was used to identify differentially expressed genes (DEGs) between the clusters. A cluster resolution of 0.4 was set as the active identity and comprised 30 clusters to be annotated. An automated annotation tool known as SCSA (Cao, Wang and Peng, 2020) was used to obtain putative cluster annotations by comparing the list of DEGs per cluster obtained from FindAllMarkers() to reference datasets of known cell-type-specific markers including SCSA’s built-in reference databases (Zhang *et al*., 2019) as well as user-defined databases curated from mousebrain.org (Zeisel *et al*., 2018). Manual inspection methods involved visualizing the expression of a smaller set of known cell-type-specific markers across the different clusters in bubbleplots as well as searching for the cell-type-specific markers in the list of cluster-specific DEGs from the FindAllMarkers() output. Using the automated and manual annotation method, putative user-defined annotations for each cluster were generated. Additionally, Seurat’s label transfer method was carried out using the Allen Mouse Brain atlas as the reference (Yao *et al*., 2021). Seurat’s FindTransferAnchors() function was used to find anchors between the reference and query datasets with “SCT” specified as the normalization method and the top 30 dimensions used. A relatively stringent filtering step was then performed to remove any nuclei that did not have agreement between their broad user-defined and Allen Mouse Brain annotations. Following this filtering step, the user-defined coarse annotations were used for downstream analyses. The 4 control samples were subsetted and heatmaps were plotted to visualize the normalized average expression of several genes of interest across the pyramidal neuron and astrocyte clusters respectively.

### Data analysis and statistics

Data analysis was performed using custom scripts written in MATLAB (MathWorks). The image analysis pipeline extracted signals from a region of interest (ROI) targeted to the soma of the cell and calculated an integrated density of fluorescence (i.e., mean pixel brightness) in this region (Fs). Background subtraction was performed by selecting a ROI targeted to a representative area of the background, calculating an integrated density of fluorescence in this region, and subtracting this from Fs image-wise. For ORCHID recordings this background fluorescence signal was smoothed with a smoothing window of 100 before being subtracted from Fs. Correction for photobleaching and low-frequency changes in signal was performed by fitting and subtracting a 9^th^- or 10^th^-order polynomial to and from the background-corrected Fs trace. The response to the GABA puff or optogenetic stimulation in the Fs trace was excluded from the fitting of this polynomial to avoid removing the signal from the trace. For recordings using Voltron2549-ST with GABAAR activation this corrected Fs trace was smoothed with a smoothing window of 7, and ΔF/F0 values were calculated using the maximum or minimum value of the smoothed signal post-puff and the mean of the smoothed baseline period prior to the puff. This same process was used for ORCHID recordings during activity-dependent variation in DFGABAA experiments, with the maximum or minimum value of the smoothed signal during an optogenetic stimulation being used to calculate ΔF/F0. For all other *in vitro* ORCHID recordings, the 5 strobed optogenetic stimulation periods in the Fs trace were averaged before ΔF/F0 was calculated using the mean during averaged optogenetic stimulation and the mean during the averaged baseline period prior to optogenetic stimulation onset. Due to the higher levels of spontaneous activity *in vivo*, for *in vivo* ORCHID recordings ΔF and F0 were calculated using the mean fluorescence during a single optogenetic stimulus and an equivalent duration of recording on one side of the stimulus (these values termed ΔFS and F0S). To correct for neuropil contamination, the indices of these two periods (the period during the stimulus and the baseline period) were used to calculate an equivalent ΔF from the background fluorescence trace (ΔFBG), which was subtracted from ΔFS (giving ΔFS-BG). ΔF/F0 was then calculated using ΔFS-BG and F0S. Using the linear fluorescence changes recorded in response to voltage steps (**Fig. 1i,j** and **Supplementary Figure 3c,d**), estimated DFGABAA was calculated from ΔF/F0. Statistical measurements were performed using GraphPad Prism. The Shapiro-Wilk test was used to test for normality, and parametric or non-parametric tests were selected as required. Data is reported as mean ± SEM unless stated otherwise.

### Computational Model

We simulated a neuron using the pump-leak mechanism as a single compartment within an extracellular environment consisting of fixed ion concentrations (“infinite bath model”). The compartment was modelled as a cylinder with a diameter of 1 µm and a length of 25 µm. Ionic flow between the compartment and external environment was permitted via leak channels, a Na^+^/K^+^-ATPase, and KCC2 transporters. For details of the simulation equations for ionic flux and volume change, see Düsterwald *et al*. (2018). In addition, GABAARs, HCO3- and pH regulation were added to the model in Düsterwald *et al*. (2018), which we describe below. Fast GABAergic synaptic transmission mediated by GABAARs was modelled as an alpha function with a tau (*τ)* of 250 ms, and a max conductance gGABAA_max of 10 nS:

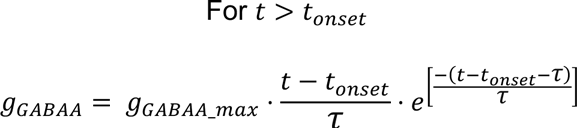

Current through GABAARs (*I_GABAA_*) was modelled as follows:

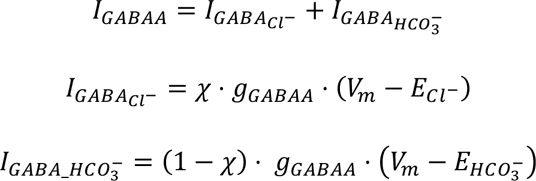

χ represents the fraction of the total GABAergic current carried by Cl^-^ and is given by:

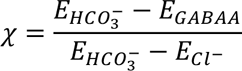

where the reversal potential for Cl^-^ (ECl-), HCO3-(EHCO3-)and the GABAA receptor (EGABAA) were updated, throughout the simulation using the Nernst and Goldman-Hodgkin-Katz equations, respectively:

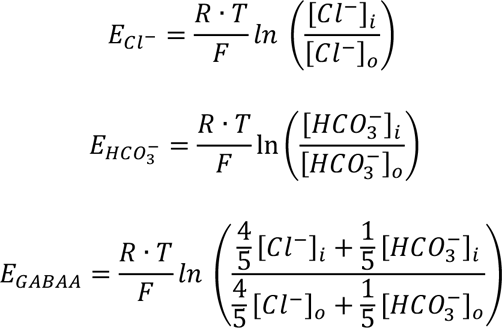

where R is the ideal gas constant, F is Faraday’s constant, and T is temperature.

Carbon dioxide was assumed to be equilibrium distributed across the simulated plasma membrane. For a pCO2 of 38 mmHg (5% CO2), [CO2] was calculated using Henry’s Law and a Henry’s Law constant for CO2 in water of 0.031 (M/atm).

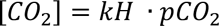

Assuming that [CO2! # $%2CO3], $%2CO3] (both inside and outside) was calculated as 1.55 mM. The CO2 hydration reaction outside the compartment was assumed to be under equilibrium.

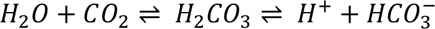

Using the Henderson Hasselbalch equation and a pKa of 6.1 as the dissociation constant for H2CO3, [HCO ^-^] was calculated at pH of 7.4 (outside):

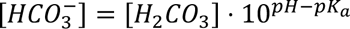

giving a [HCO ^-^] = 31 mM.

Inside the compartment, pH was held constant at 7.2 and the production HCO3-by the forward reaction of the CO2 hydration reaction was calculated using the forward rate equation and a forward rate constant Kf of 1×10^3^:

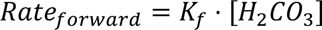

Whilst the removal of HCO3-by the reverse reaction was calculated using the reverse rate equation and a reverse rate constant Kr of 2.539×10^9^:

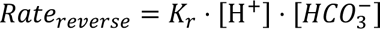

H+ produced or removed by the forward or reverse reactions was simulated as being immediately exchanged with Na^+^ via the Na^+^/H^+^ exchanger (Chesler, 2003) in order to maintain an intracellular pH at 7.2.

Parameters:

| Symbol | Value | Description |
| --- | --- | --- |
| **Constants** |  |  |
| F | 96485.33 C/mol | Faraday constant |
| R | 8.31446 J/(K·mol) | Universal gas constant |
| T | 310.15 K | Absolute temperature (= 37°C) |
| **Parameters** |  |  |
| Radius | 5 µm | Radius of the compartment |
| Length | 25 µm | Length of the compartment |
| C<sub>m</sub> | 2 x10<sup>-6</sup> F/cm<sup>2</sup> | Unit membrane capacitance (Qian and Sejnowski, 1989) |
| $g_{Na}$ | 25 $\mu S/cm^2$ | $Na^+$ leak conductance |
| $g_K$ | 70 $\mu S/cm^2$ | $K^+$ leak conductance |
| $g_{Cl}$ | 20 $\mu S/cm^2$ | $Cl^-$ leak conductance |
| $g_{HCO_3}$ | 4 $\mu S/cm^2$ | $HCO_3^-$ leak conductance |
| $g_{KCC2}$ | 20 $\mu S/cm^2$ | KCC2 conductance |
| $v_w$ | 0.018 $dm^3/mol$ | Partial molar volume of water<br>(Hernández and Cristina, 1998) |
| $p_w$ | 0.0015 $dm/s$ | Osmotic permeability (Hernández and Cristina, 1998) |
| $P$ | 0.1 $C/(dm^2 \cdot s)$ | ATPase pump rate constant |
| $pCO_2$ | 0.05 atm | Partial pressure of $CO_2$ |
| $k_H$ | 0.031 $M/atm$ | Henry's law constant for $CO_2$ in water |
| $K_f$ | $10^3 s^{-1}$ | Forward rate constant |
| $K_r$ | $2.539 \times 10^9 s^{-1} \cdot M^{-1}$ | Reverse rate constant |
| $g_{GABAA\_max}$ | 10 nS | Maximum conductance of $GABA_A$Rs. |
| $pH_o$ | 7.4 | Extracellular pH |
| $pH_i$ | 7.2 | Intracellular pH |
| $[K^+]_o$ | 3.5 mM | Extracellular $K^+$ conc. |
| $[Na^+]_o$ | 145 mM | Extracellular $Na^+$ conc. |
| $[Cl^-]_o$ | 110 mM | Extracellular $Cl^-$ conc. |
| $[HCO_3^-]_o$ | 31 mM | Extracellular $HCO_3^-$ conc. |
| $[H_2CO_3]_o$ | 1.55 mM | Extracellular $H_2CO_3$ conc. |
| $[H_2CO_3]_i$ | 1.55 mM | Intracellular $H_2CO_3$ conc. |
| $[X^-]_o$ | 7.5 mM | Extracellular impermeant anion conc. |
| $z_o$ | -1 | Extracellular impermeant anion average charge |
| $z_i$ | -0.85 | Intracellular impermeant anion average charge (Lodish et al. 2009) |

| Variables (default steady state) |  |  |
| --- | --- | --- |
| $[K^+]_i$ | 122.6 mM | Initial intracellular $K^+$ conc. |
| $[Na^+]_i$ | 15 mM | Initial intracellular $Na^+$ conc. |
| $[Cl^-]_i$ | 5.3 mM | Initial intracellular $Cl^-$ conc. |
| $[HCO_3^-]_i$ | 9.7 mM | Initial intracellular $HCO_3^-$ conc. |
| $[X^-]_i$ | 144.4 mM | Initial intracellular impermeant anion conc. |
| $V_m$ | -67.2 mV | Initial membrane potential |
| $w$ | 19 pL | Compartment volume |

Code used for the simulations is accessible at: https://github.com/Eran707/Single-Cell-Simulator.

## Data and code availability

All data was analyzed using custom scripts created in Matlab. Code will be made publically available on GitHub upon publication, but until then is available upon request. Source data will be published alongside the publication of the manuscript and raw data is available upon reasonable request.

## Acknowledgements

We would like to thank members of the Raimondo lab for advice and comments and to Suzanne Duncan with assistance naming ORCHID. We thank Sarah E. Plutkis, Katie L. Holland, Jonathan B. Grimm, and Luke D. Lavis (Janelia) for Janelia Fluor HaloTag ligands and Yoav Adam for providing advice on GEVI imaging. We thank Rodney Lucas for assistance performing animal experiments and Christina Steyn and Dorit Hockman for advice performing snRNAseq analysis. The research leading to these results has received support from the Gabriel Foundation, a Wellcome Trust Seed Award (214042/Z/18/Z), the FLAIR Fellowship Programme (FLR\R1\190829): a partnership between the African Academy of Sciences and the Royal Society funded by the UK Government’s Global Challenges Research Fund, a Wellcome Trust International Intermediate Fellowship (222968/Z/21/Z), ERC Grant Agreement 617670 and the project was supported by funding from BBSRC project BB/S007938/1.

## Author contributions

Conceptualization: JSS, ASA, ERS, CJA, JVR

Tool provision: ASA, ERS

Investigation: JSS, TJSS, RJB, EFS, KMD, SEN, JVR

Writing: JSS, CJA, JVR Supervision: JVR

Funding acquisition: CJA, JVR

## Supplemental information

**Supplementary Figure 1.**
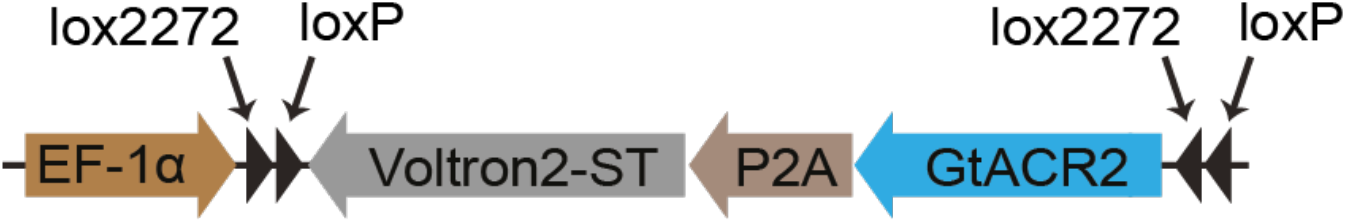
The ORCHID construct. Voltron2-ST is expressed together with GtACR2 via a P2A linker sequence, with a Flex design and expression under the EF-1α promoter.

**Supplementary Figure 2:**
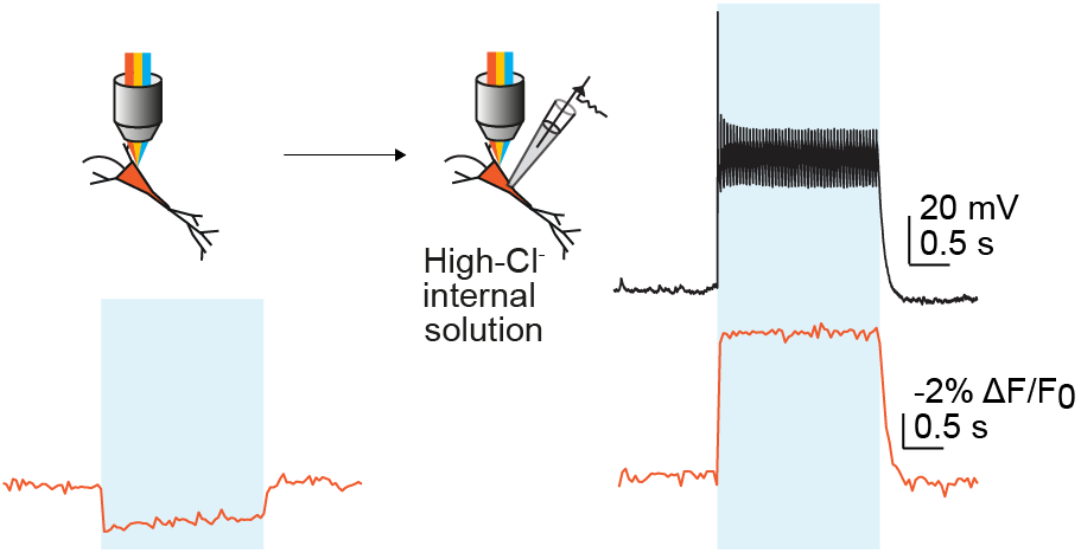
ORCHID can detect changes in DFGABAA induced by an increase in [Cl^-^]i imposed by dialysis. Schematic (top left) showing ORCHID used to record DFGABAA in a neuron at baseline (bottom left, orange trace) and again after being patched with high-Cl^-^ (141 mM) intracellular solution (right). Both voltage imaging (bottom right, orange trace) and current-clamp recordings (top right, black trace) showed a transition to a strongly depolarizing DFGABAA. Action potentials are truncated due to the low-pass behavior of the image acquisition frequency (25 Hz) and averaging of responses (see **Methods**).

**Supplementary Figure 3:**
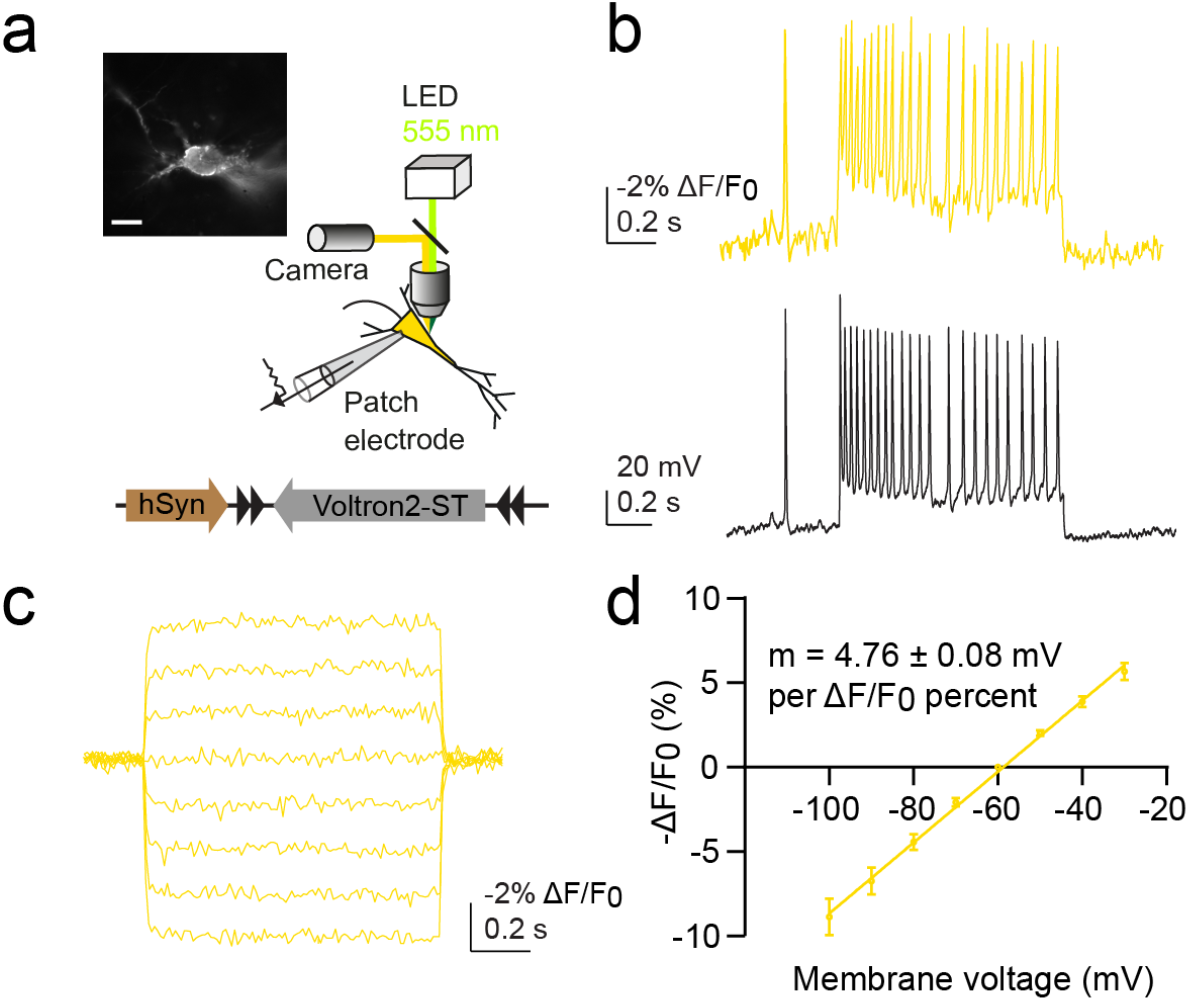
Characterizing Voltron2549-ST for measurement of Vm. **a**, Schematic of experimental setup for patch-clamp characterization of Voltron2549-ST in mouse hippocampal organotypic brain slice cultures. Inset: widefield fluorescence image of the neuron patched in ‘b’; scale bar: 10 µm. **b**, Current-clamp recording (black trace) and the Voltron2549-ST fluorescence response (yellow trace) following a 200 pA current injection. **c**, Voltron2549-ST fluorescence response to 10 mV voltage steps (Vhold = -60 mV). **d**, Voltage-fluorescence relation of Voltron2549-ST, with a slope 4.76 ± 0.08 mV per ΔF/F0 percent (R^2^ = 0.9236, *n* = 7).

**Supplementary Figure 4:**
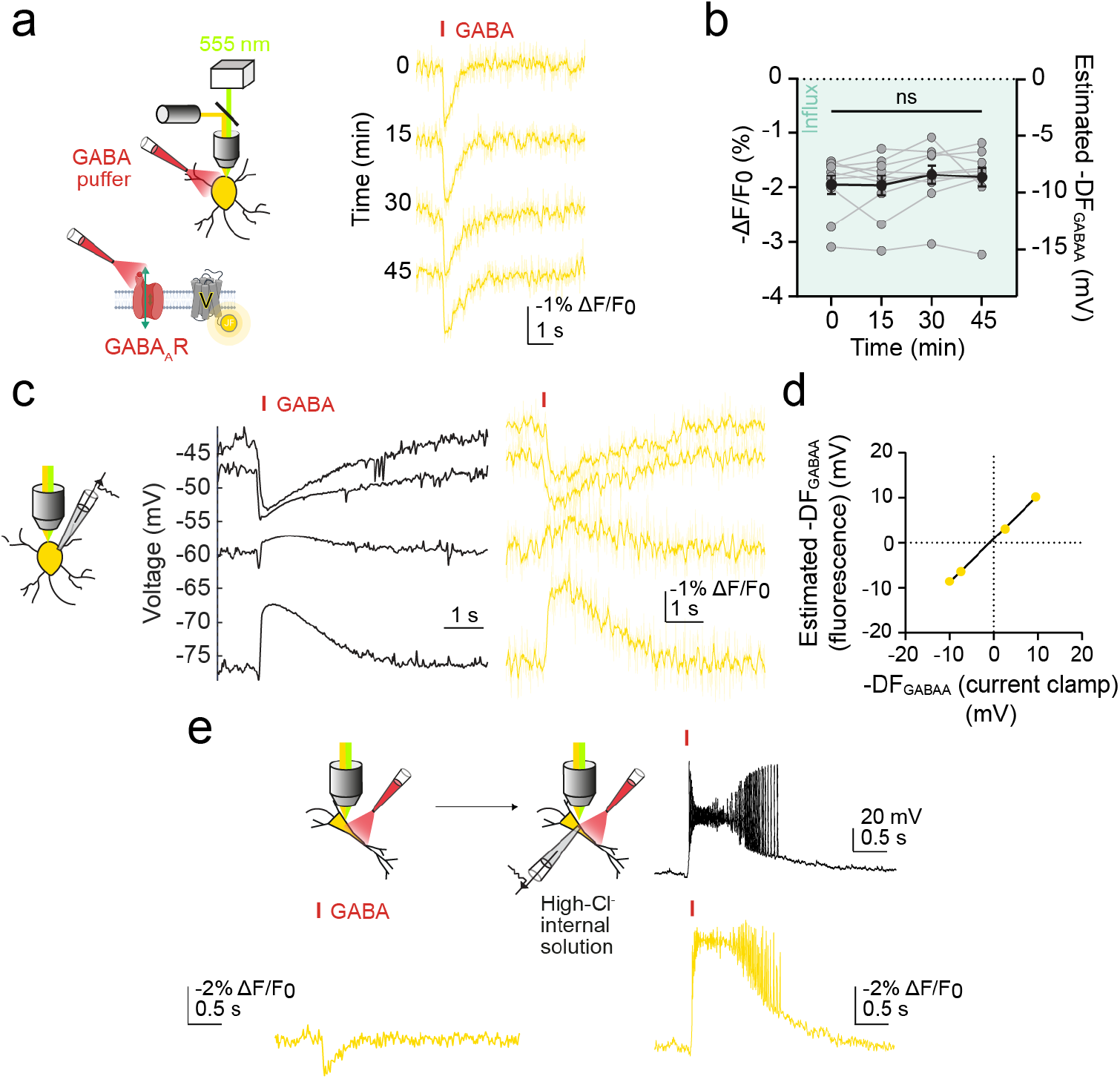
Characterizing Voltron2549-ST for measurement of DFGABAA via activation of endogenous GABAARs. **a**, Left, schematic demonstrating imaging of Voltron2549-ST together with activation of endogenously expressed GABAARs through picolitre delivery of 500 µM GABA (red) directed at the cell soma, with 5 µM CGP-55845 used to block GABABRs. Right, repeated fluorescence transients (yellow traces) in response to GABA application (red bar) recorded over 45 min. **b**, Population data showing estimates of DFGABAA calculated from ΔF/F0 remained stable (Friedman test, *P* = 0.2933, *n* = 10). Green shading indicates direction of estimated anion flux. **c**, Voltron2549-ST and activation of GABAARs were used to estimate DFGABAA in neurons current-clamped at a range of Vm values (left panel, Vm; right panel, fluorescence). **d**, The voltage-fluorescence relation of Voltron2549-ST (**Supplementary Fig. 3d**) was used to convert ΔF/F0 measurements at each Vm to estimated DFGABAA values. These were equivalent to the DFGABAA values recorded at each Vm using the current-clamp recording (R^2^ = 0.9996). **e**, Schematic (top left) showing Voltron2549-ST and activation of GABAARs used to estimate DFGABAA in neurons at baseline (bottom left, yellow trace) and again after being patched with high-Cl^-^ (141 mM) intracellular solution. Both voltage imaging (bottom right, yellow trace) and current-clamp recordings (top right, black trace) showed a transition to a strongly depolarizing response to GABAAR activation after Cl^-^ loading. ns = not significant (*P* > 0.05); error bars indicate mean ± SEM.

**Supplementary Figure 5:**
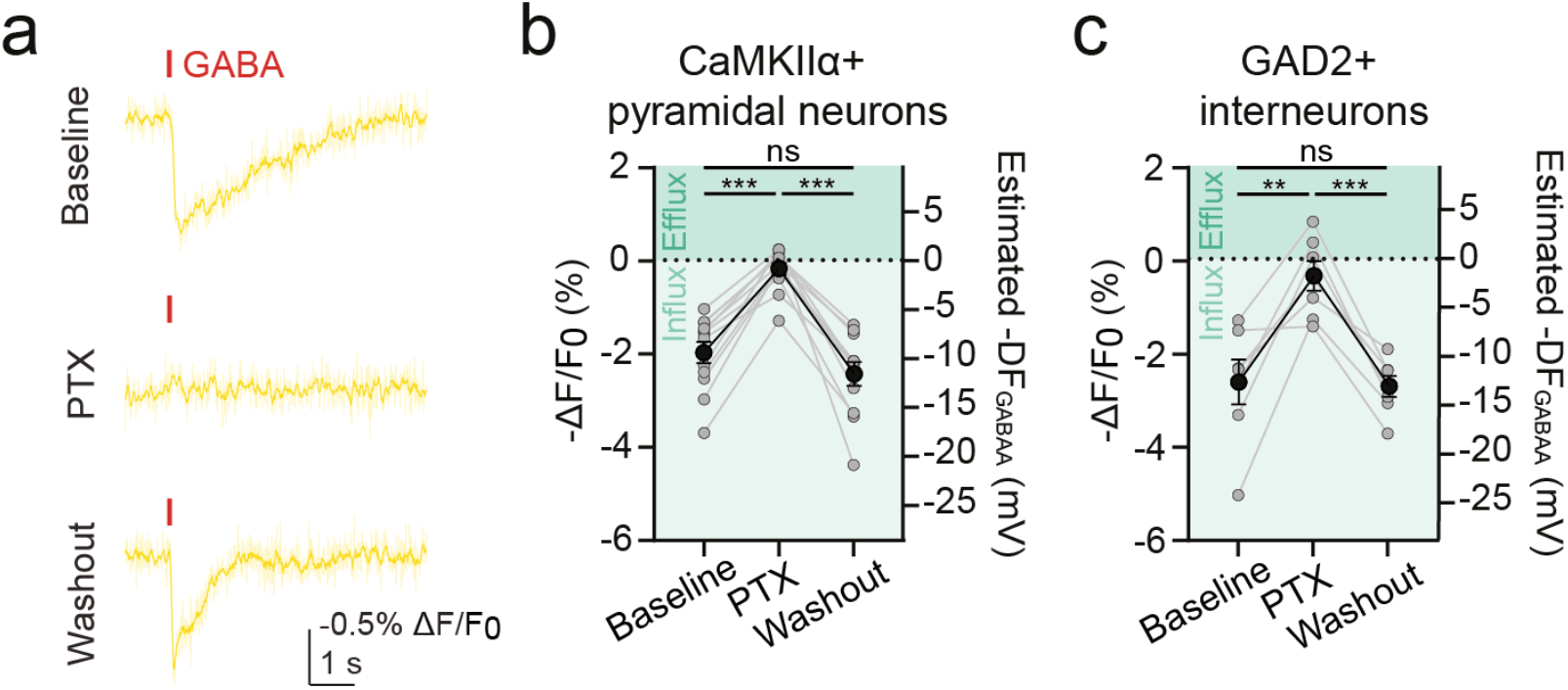
Voltron2549-ST estimates of agonist-evoked DFGABAA reflect GABAAR activity. **a**, Voltron2549-ST and GABA puffs were used to record DFGABAA in virally transfected organotypic hippocampal neurons at baseline, during picrotoxin (PTX; 100 µM) wash-in, and again after PTX washout. **b**, Population data showing recordings of DFGABAA in CaMKIIα+ pyramidal neurons measured hyperpolarizing DFGABAA at baseline, with this response disappearing upon PTX wash-in (Wilcoxon matched-pairs signed rank test, *P* = 0.0005, *n* = 12). The hyperpolarizing DFGABAA response returned upon PTX washout (baseline vs washout: paired t-test, *P* = 0.1256, *n* = 12; PTX vs washout: Wilcoxon matched-pairs signed rank test, *P* = 0.0005, *n* = 12). **c**, Population data for GAD2+ interneurons (baseline vs PTX: paired t-test, *P* = 0.0032, *n* = 7; baseline vs washout: paired t-test, *P* = 0.8604, *n* = 7; PTX vs washout: paired t-test, *P* = 0.0002, *n* = 7). Green shading indicates direction of estimated Cl^-^ flux. ns = not significant (*P* > 0.05); \*\**P* ≤ 0.01; \*\*\**P* ≤ 0.001; error bars indicate mean ± SEM.

**Supplementary Figure 6:**
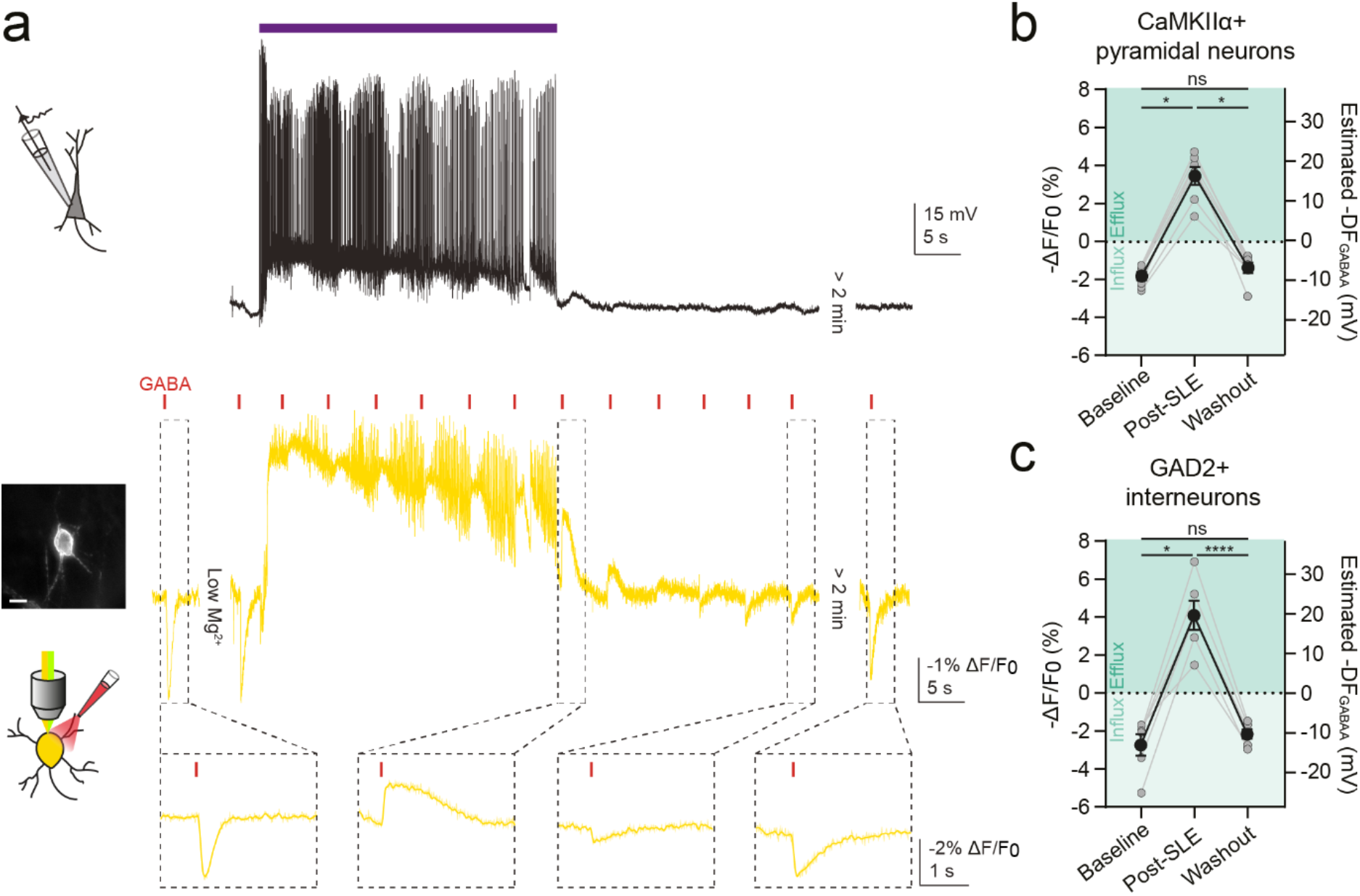
Cell-type-specific dynamics in DFGABAA recorded using Voltron2549-ST and GABAAR activation during SLEs. **a**, Voltron2549-ST and GABAAR activation were used to investigate activity-dependent variation in DFGABAA in neurons in mouse hippocampal organotypic brain slice cultures. SLEs were induced using the low-Mg^2+^ model of *in vitro* SLEs. A whole-cell current-clamp recording from a CA1/3 pyramidal neuron provided an independent readout of SLEs (top panel; purple bar indicates SLE). Concurrent Voltron2549-ST recordings were used to simultaneously record DFGABAA in target cells, whilst also providing a recording of the SLE (bottom panel, yellow traces). Central inset image: widefield fluorescence image of the GAD2+ interneuron from which the recording was made; scale bar: 10 µm. Baseline recordings of DFGABAA were made prior to low-Mg^2+^ aCSF wash-in (labelled as “Low Mg^2+^”), immediately post-SLE (< 20 seconds after SLE cessation), and > 2 min after SLE cessation (washout). **b**, Population data of DFGABAA in CaMKIIα+ pyramidal neurons showed a shift from hyperpolarizing DFGABAA at baseline to depolarizing DFGABAA post-SLE (Wilcoxon matched-pairs signed rank test, *P* = 0.0156, *n* = 6). > 2 min after the SLE (washout), DFGABAA had returned to baseline levels (baseline vs washout: Wilcoxon matched-pairs signed rank test, *P* = 0.1563, *n* = 7; post-SLE vs washout: Wilcoxon matched-pairs signed rank test, *P* = 0.0156, *n* = 7). **c**, Similar results were recorded in GAD2+ interneurons (baseline vs post-SLE: Wilcoxon matched-pairs signed rank test, *P* = 0.0313, *n* = 7; baseline vs washout: Wilcoxon matched-pairs signed rank test, *P* = 0.2188, *n* = 6; post-SLE vs washout: paired t-test, *P* < 0.0001, *n* = 6). Green shading indicates direction of anion flux. ns = not significant (*P* > 0.05); \**P* ≤ 0.05; \*\**P* ≤ 0.01; \*\*\*\**P* ≤ 0.0001; error bars indicate mean ± SEM.

**Supplementary Figure 7:**
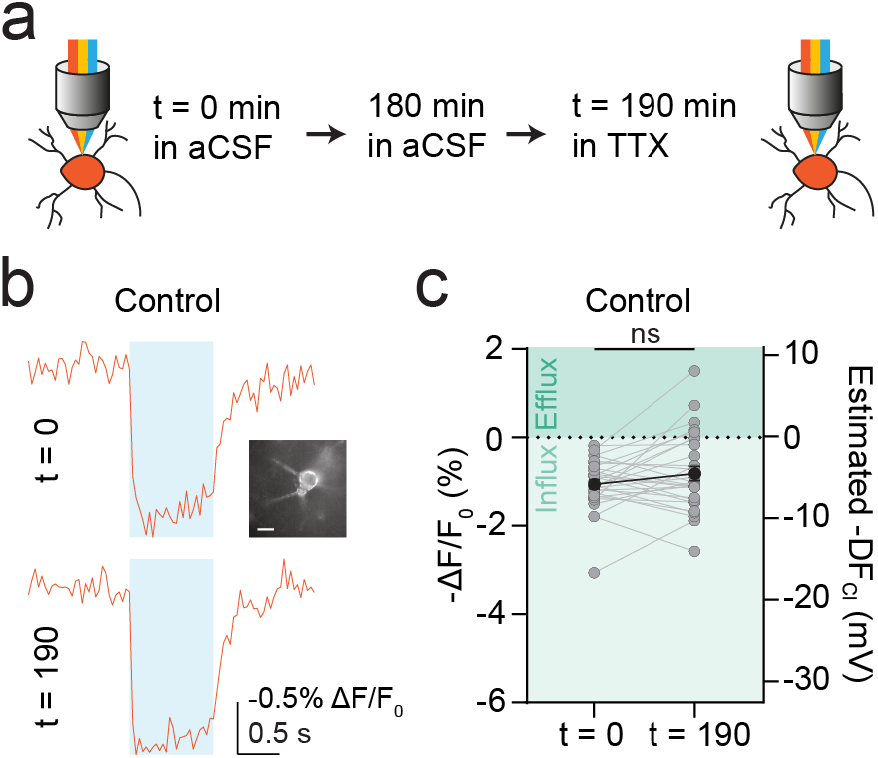
ORCHID estimates of DFGABAA are stable over 190 min. **a**, Schematic of experimental design with ORCHID used to make paired recordings of DFGABAA 190 min apart in the same GAD2+ interneurons in organotypic hippocampal brain slices. **b**, Top, baseline recording from a GAD2+ interneuron before slice was incubated in normal aCSF for 180 min. Bottom, another recording from the same cell after the slice had been transferred to aCSF containing TTX (1 µm) for 10 min. Inset: widefield fluorescence image of recorded neuron; scale bar = 10 µm. **c**, Population data shows DFGABAA was not significantly different at t = 0 and t = 190 (Wilcoxon matched-pairs signed rank test, *P* = 0.2297, *n* = 29). ns = not significant (*P* > 0.05); error bars indicate mean ± SEM.

